# Apolipoprotein E governs neuroinflammation resolution and recovery from viral encephalitis

**DOI:** 10.64898/2026.04.28.721524

**Authors:** Abigail L. Cox, Wilson Nguyen, Bing Tang, Kexin Yan, Ashwini Raghavendra, Rebekah Ziegman, Gunter Hartel, Quan H. Nguyen, Mark P. Hodson, Andreas Suhrbier, Cameron R. Bishop, Daniel J. Rawle

## Abstract

Effective control of viral encephalitis requires immune responses that clear infection without causing damaging neuroinflammation, yet the mechanisms governing resolution and recovery remain unclear. Using a Japanese encephalitis virus mouse model spanning asymptomatic, symptomatic, and lethal trajectories, together with single-cell spatial transcriptomics and RNA-seq, we identify apolipoprotein E (ApoE) as a driver of neuroinflammation resolution. *Apoe* was upregulated in microglia and infiltrating myeloid cells of symptomatic survivors, where *Apoe-Trem2-Tyrobp* signalling promoted a phagocytic, anti-inflammatory program. In contrast, immune cells in lethal disease failed to induce *Apoe* and remained pro-inflammatory. ApoE-deficient mice were unable to recover following encephalitis onset, demonstrating that ApoE signalling is essential for resolution and recovery. Analysis of cerebrospinal fluid from acute encephalitis patients linked APOE isoforms to neuroinflammation resolution, with APOEε2 carriers exhibiting reduced neutrophils and shorter hospitalisation. These findings identify ApoE as a critical driver of neuroinflammation resolution and a promising therapeutic target for viral encephalitis.

## INTRODUCTION

Japanese encephalitis virus (JEV) is a neurotropic, single-stranded positive-sense RNA virus of the *Orthoflavivirus* genus. JEV circulates in enzootic cycles involving *Culex* mosquitoes and vertebrate hosts such as pigs and wading birds^1^, with humans considered dead-end hosts because viremia is typically too low to infect feeding mosquitoes^2,3^. Although most human infections are asymptomatic, approximately 1 in 250 progresses to encephalitis. These encephalitic cases carry a high fatality rate of 20-30%, while the survivors frequently experience long-term neurological sequelae, including seizures, motor impairments, and cognitive or speech dysfunction^4^. JEV is a leading cause of viral encephalitis across Asia and the Western Pacific, resulting in an estimated 70,000 clinical cases each year^4^. Despite the availability of effective vaccines^5^, the substantial disease burden and high mortality among symptomatic cases underscore the pressing need for improved understanding of JEV neuropathogenesis and the development of targeted therapeutics.

After transmission by a mosquito bite, JEV initially replicates in peripheral myeloid cells, generating a systemic viremia^6^. As viremia develops, JEV infection activates perivascular mast cells at the blood-brain barrier (BBB), leading to the release of chymase which cleaves tight junction proteins and enhances BBB disruption and viral neuropenetration^7^. Once inside the brain, JEV replicates in perivascular glial populations (including oligodendrocytes and oligodendrocyte progenitor cells), triggering a strong type I interferon (IFN) response in neighbouring uninfected glial cells^8^, which restrains *orthoflavivirus* replication in these cells^9^. Uncontrolled viral replication in neurons^10–12^ activates pro-inflammatory pathways, driving the influx of infiltrating immune cells such as monocytes, macrophages, T cells, and natural killer cells, and activating resident astrocytes and microglia^12–14^. The resulting inflammatory cascade leads to high levels of pro-inflammatory cytokines and chemokines, including TNF-α, IL-6, IL-1β, and CCL2^12,15^, ultimately contributing to neuronal injury and cell death^11,12,15,16^. In mice and non-human primate models, as well as in humans, this encephalitis is often fatal. However, some individuals and animals can successfully clear JEV from the brain and the encephalitis can resolve without causing death or permanent neurological sequelae. In general, the resolution of inflammation relies on coordinated immune mechanisms that restore tissue homeostasis^17^. Pro-resolving mediators shift the immune response from a pro- to an anti-inflammatory state, gradually reducing cytokine and chemokine levels. This phase is marked by changes in immune-cell phenotypes, including a shift from Th1 to Th2 cells and Tregs, and from pro-inflammatory, antiviral M1 macrophages to anti-inflammatory, tissue-repairing M2 macrophages^17^. Despite extensive research into *orthoflavivirus* neuroinvasion and neuropathogenesis, the molecular mechanisms governing the resolution of neuroinflammation and the re-establishment of homeostasis after *orthoflavivirus* encephalitis remain completely undefined.

Human brain samples are rarely available at the critical junctures when neuroinflammation either escalates or resolves. As a result, immunocompetent laboratory mouse models have become essential for studying the neuroinvasion and neuropathogenesis of lethal JE, and they successfully recapitulate many key features of human and primate disease^11,18^. When combined with recent advances in single cell and spatially resolved transcriptomics, these mouse models have substantially advanced our understanding of JEV neuroinvasion and early innate immune responses^8^ and the mechanisms driving pathogenic neuroinflammation^13,19^. However, mouse models have not been used to investigate disease resolution following *orthoflavivirus* encephalitis, hampering our understanding of this process and the development of disease resolving therapies.

Here, we used a mouse model of Japanese encephalitis that recapitulates asymptomatic, symptomatic, and lethal disease outcomes^11^. Using this model, we show that *Apoe* is markedly upregulated in brain immune cells of symptomatic mice that survive infection, but not in mice with lethal outcomes. These *Apoe*-expressing immune cells signal via Trem2-Tyrobp and adopt a phagocytic, anti-inflammatory phenotype. In contrast, ApoE-deficient mice exhibit dysregulated immune responses, more severe disease, and an inability to recover once symptomatic, demonstrating that ApoE is essential for successful resolution of JEV neuroinflammation. In addition, we show that APOE isoform impacts markers of inflammation and recovery from infectious acute encephalitis syndrome (AES) in a human cohort. Collectively, our findings identify ApoE upregulation in brain immune cells as a central orchestrator of pro-resolution immune programs that restoration of homeostasis and survival following viral encephalitis.

## MATERIALS AND METHODS

### Ethics statement and regulatory compliance

All mouse work was conducted in accordance with the “Australian code for the care and use of animals for scientific purposes” as defined by the National Health and Medical Research Council of Australia. Mouse work was approved by the QIMR Berghofer Animal Ethics Committee (P3746, A2108-612). Mice were euthanised using CO_2_. Breeding and use of GM mice was approved under a Notifiable Low Risk Dealing (NLRD) Identifier: NLRD_Suhrbier_Oct2020: NLRD 1.1(a). All infectious JEV work was conducted in a dedicated suite in a biosafety level-3 (PC3) facility at QIMR Berghofer (Australian Department of Agriculture, Water and the Environment certification Q2326 and Office of the Gene Technology Regulator certification 3445). All work was approved by the QIMR Berghofer Safety Committee (P3746). Research with JEV were approved under the Queensland Biosecurity Act, Scientific Research Permit (restricted matter) – Permit number PRID000916.

### Single-cell spatial transcriptomics slide preparation

Formalin fixed C57BL/6J brains were embedded in paraffin and sections were prepared using the CosMx™ Spatial Molecular Imager (SMI) Sample Preparation kit (nanoString, Bruker Spatial Biology, Inc, USA) as per manufacturers’ instructions. Histopathological analyses of these brains were reported previously^11^. The run included 2 uninfected, 2 JEV_Nakayama_ lethal, 1 JEV_FU_ lethal, 1 MVEV_TC123130_ lethal, and 1 JEV_FU_ symptomatic recovered day 30 post infection across two CosMx slides. A mouse neuroscience panel of RNA probes consisting of 950 mRNA targets and custom targets for JEV_FU_, JEV_Nakayama_, and MVEV_TC123130_ (Supplementary Table 1). The prepared slides were loaded onto the CosMx™ SMI and using the CosMx™ SMI Control Center, ∼200 fields of view were chosen for each slide. Fields of view were selected based on the presence of viral infection and JEV-associated brain lesions, as identified by H&E and 4G4 staining previously^11^, and were also chosen to capture representative areas of each brain region consistently across samples.

### Single-cell spatial transcriptomics analysis pre-processing

Quality control (QC) metrics were generated using the AtoMx Spatial Informatics Platform (NanoString). Data were exported from AtoMx and further analysed in R v4.2 using Seurat v5.0 and third-party packages where noted. Cells with less than five gene counts were removed, and negative/system-control probes were excluded. Per-slide Seurat objects were merged into a single object. To identify cell type clusters, the merged object was integrated using Seurat’s reciprocal principal components analysis (RPCA) method, followed by normalisation, variable feature identification, scaling, and principal component analysis (PCA) with 25 PCs. A nearest-neighbour graph was built, uniform manifold approximation and projection (UMAP) was computed, and clustering was performed using the FindClusters function with resolutions of 0.1, 0.2, 0.3 and 0.4. The Clustree package was used to examine clustering robustness.

Transcript coordinates, cell boundaries, and cluster identities (at resolution 0.2) were used in combination with the FastReseg package to evaluate segmentation errors and refine cell boundaries and per-cell transcript counts. Updated transcript counts were used to reconstruct a new Seurat object, which was pre-processed as above, with clustering performed at resolutions 0.01, 0.05, 0.2, and 0.5. Using 0.5 resolution clusters, a further round of sub-clustering was performed within each of the T cell, astrocyte, and microglia clusters using a resolution of 0.1. Cell-type clusters were annotated manually using a combination of the topmost differentially expressed genes identified by the FindAllMarkers function in Seurat, and canonical marker genes obtained from the Allen Brain Atlas and selected publications^20,21^ (Supplementary Table 1). Marker gene expression was examined using the scCustomize package.

### Pseudo-bulk differential expression analysis

Bootstrapped pseudo-bulk differential expression analysis was performed by creating a count matrix of pseudo-samples, in which each pseudo-sample was the summation of counts for cells related to a given section/cell-type combination. Pseudo-samples with ≥200 cells were analysed. 100 bootstrap iterations (n=200 cells each iteration, with replacement) were averaged to produce final pseudo-counts. Differential expression analysis was performed using DESeq2, with a significance threshold of FDR-adjusted p < 0.05. Normalised, variance-stabilised transformed (vst) counts were used for PCA and heatmaps. DEG sets were analysed with Ingenuity Pathway Analysis (Qiagen) Core Analysis tool, and the gprofiler2 package using the Gene Ontology, KEGG, and Reactome databases.

### Receptor-ligand interaction analysis

Paracrine signalling was investigated using the spatialDM package in Python with parameters length_scale = 25 µm; distance_cutoff = 0.1; n_neighbors_broad = 80; n_neighbors_tight = 6; single_cell = True. The ‘single_cell = True’ parameter excludes autocrine signalling from the analysis. For reference, the CellChatDB v2 database^22^ was combined with the following set of *Apoe* ligand receptor interactions: Apoe-Chrna4, Apoe-Lrp2, Apoe-Lrp6, Apoe-Lsr, Apoe-Sorl1, and Apoe-Trem2 from CellTalk_Mm v1^23^, and Apoe-Tyrobp from CellPhone v5^24^. Results were examined and plotted using the Seurat, circlize and pheatmap packages in R v4.4.2. Cell types most involved in *Apoe* signalling (per SpatialDM) were analyzed for differential expression by defining ‘*Apoe*-receiver’ cells as those with SpatialDM p < 0.05 and receptor Moran’s I > 1. Then, for each cell-type of interest, a set of ‘*Apoe*-non-receiver’ cells was created by identifying the nearest unique neighbors of cells in the *Apoe*-receiver set that were not identified by SpatialDM as being involved in any *Apoe* interaction. Differentially expressed genes were identified using the FindMarkers function from the Seurat package with parameters: logfc.threshold=0.25; min.pct=0.1; test.use=’t’. *Apoe* was excluded from differential expression testing.

### Cell lines and culture

Vero E6 (C1008, ECACC, Wiltshire, England; obtained via Sigma Aldrich, St. Louis, MO, USA) and C6/36 cells (ATCC# CRL-1660) were cultured in medium comprising RPMI 1640 (Gibco), supplemented with 10% foetal bovine serum (FBS), penicillin (100 IU/ml)/streptomycin (100 μg/ml) (Gibco/Life Technologies) and L-glutamine (2 mM) (Life Technologies). Vero E6 cells were cultured at 37°C and 5% CO_2_, and C6/36 cells were cultured at 28°C and 5% CO_2_.

### Virus isolates and culture

JEV_FU_ (GenBank: AF217620) was obtained from A/Prof. Gregor Devine (QIMR Berghofer). Virus stocks were generated by infection of C6/36 cells at multiplicity of infection (MOI) ≈0.1, with supernatant collected after ∼5 days, cell debris removed by centrifugation at 3000 × g for 15 min at 4°C, and virus aliquoted and stored at -80°C. Virus stocks used in these experiments underwent less than three passages in our laboratory. Virus titres were determined using standard CCID_50_ assays (see below). Virus stocks were sequenced and compared with the reference genome using Integrative Genomics Viewer (v2.11.2) which confirmed the identity of the virus stock as JEV_FU_.

### Cell Culture Infectious Dose 50% (CCID_50_) assays

CCID_50_ assays were performed as previously described^25–27^. C6/36 cells were plated in 96 well flat bottom plates at 2×10^4^ cells per well in 100 µl of medium. Samples underwent 10-fold serial dilutions in 100 µl RPMI 1640 supplemented with 2% FBS, performed in duplicate for serum and cell culture supernatant. For serum, a volume of 100 µl of serially diluted samples were added to each well of 96 well plates containing C6/36 cells, and the plates were cultured for 5 days at 37°C and 5% CO_2_. 25 µl of supernatant from infected C6/36 cells were then passaged on to Vero E6 cells plated the day before at 2×10^4^ cells per well in 96 well flat bottom plates. Supernatant from *in vitro* cultures were titred directly on to Vero E6 cells. Vero E6 cells were cultured for 5 days and cytopathic effect was scored, and the virus titre was calculated using the method of Spearman and Karber^28^ (a convenient Excel CCID_50_ calculator is available at https://www.klinikum.uni-heidelberg.de/zentrum-fuer-infektiologie/molecular-virology/welcome/downloads).

### Mouse infections

C57BL/6J and *Apoe^-/-^* mice were purchased from OzGene, Bentley, WA, Australia. C57BL/6J and *Apoe^-/-^* mice used in this study were approximately 6 weeks old and were all female except for the C57BL/6J versus *Apoe^-/-^*mice experiment (Figure 6) which used an equal number of males and females for each strain. Mice were infected subcutaneously (s.c.) at the base of the tail with 100 µl of 5×10^3^ CCID_50_ virus inoculum. Serum was collected via lateral tail vein nicks into Microvette Serum-Gel 500 µl blood collection tubes (Sarstedt, Numbrecht, Germany). Mice were weighed and monitored for disease manifestations and were humanely euthanised using CO_2_ based on a previously described score card system^11^. At necropsy, the left brain hemisphere was fixed in 10% formalin for histology and single cell spatial transcriptomics, and the right brain hemisphere was stored in RNAlater for RNA-seq.

### Riluzole hydrochloride treatment of mice

Riluzole Hydrochloride (MedChemExpress) was reconstituted in dimethyl sulfoxide (DMSO) at a concentration of 100 mg/mL prior to dilution. C57BL/6J mice were treated with 200 µg/mL Riluzole Hydrochloride (+2% w/v DMSO) via drinking water. Treatment began either one day prior to infection or six days post infection and was refreshed every two days until the end of experiment. Based on the volume of water consumed in each cage, we estimate the actual dose per day per mouse to be approximately 427 µg/mouse/day, with the lower limit of 222.2 µg/mouse/day and the upper limit of 666.7 µg/mouse/day.

### Histopathology and immunohistochemistry

Formalin fixed brains were embedded in paraffin, and sections were affixed to positively charged adhesive slides and air-dried overnight at 37°C. Sections were dewaxed and rehydrated through xylol and descending graded alcohols to water. Sections were stained with H&E (Sigma Aldrich) or IHC as described below, and slides were scanned using Aperio AT Turbo (Aperio, Vista, CA USA) and analysed using Aperio ImageScope software (LeicaBiosystems, Mt Waverley, Australia) (v10). Mouse brain sections with dual OPAL immunohistochemistry staining were scanned using Zeiss Axioscan 7 fluorescence slide scanner (Zeiss, Germany) and analysed using ZEN (Blue) Microscopy software (Zeiss, Germany). Leukocyte infiltrates were quantified by measuring nuclear (strong purple staining)/cytoplasmic (total red staining) pixel ratios in scanned H&E stained images, undertaken using the Aperio Positive Pixel Count Algorithm (Leica Biosystems).

Anti-orthoflavivirus non-structural protein 1 (NS1) immunohistochemistry using 4G4^26,29,30^ was performed as explained in detail previously^11^. Anti-Ly6G (ab2557, Abcam) and anti-F4/80 (ab6640, Abcam) staining with signal development using DAB was performed as previously described^31–33^. DAB staining was quantified using the Aperio Color Deconvolution Algorithm v9.

For anti-double-stranded RNA (dsRNA) using 2G4 IHC, antigen retrieval was performed in Dako Epitope Retrieval Solution (pH 9.0) at 100°C for 20 min. Primary antibody 2G4 (mouse anti-dsRNA^34^) was diluted 1:200 in Da Vinci Green diluent and applied to the sections overnight at 4°C. The rest of the protocol followed that of 4G4^11^, except for secondary antibody was Perkin Elmer Goat anti-mouse HRP diluted 1:300 in TBST, and signal development was performed in DAB for 5 minutes after which they were washed three times in tap water.

For anti-Iba1 IHC, antigen retrieval was performed in Dako Epitope Retrieval Solution (pH 6.0) buffer and subjected to heat antigen retrieval (100°C for 20 min) using the Biocare Medical de-cloaking chamber, and slides allowed to cool for 20 minutes before transferring to TBS plus 0.025% Tween-20 (TBS-T). Non-specific antibody binding was inhibited by incubation with Biocare Medical Background Sniper with 2% BSA for 30 minutes. Rabbit anti-Iba1 (Abcam, ab178846) was diluted 1:1000 in Da Vinci Green diluent and applied to the sections for 1 hour at room temperature. Sections were washed three times in TBS-T and endogenous peroxidase was blocked by incubating slides in Biocare Medical Peroxidazed 1 for 5 minutes. Sections were washed three times in TBS-T and Perkin Elmer Goat anti-rabbit HRP diluted 1:500 in TBST was applied for 60 minutes. Sections were washed three times in TBS-T and signal developed in Vector Nova Red for 5 minutes after which they were washed three times in tap water. Sections were lightly counterstained in Haematoxylin (program 7 Leica Autostainer), washed in tap water, dehydrated through ascending graded alcohols, cleared in xylene, and mounted using DePeX or similar.

For the dual IHC staining of 4G4 and NeuN (Merck Millipore, ZMS337), endogenous peroxidase was blocked by incubating slides in Biocare Medical Peroxidazed 1 for 5 minutes. Sections were washed three times in TBS-T and goat anti-mouse Perkin Elmer Opal HRP Polymer diluted 1:500 in TBST was applied for 60 minutes. For 4G4, antigen retrieval was performed in Dako Epitope Retrieval Solution (pH 9.0) at 100°C for 20 min. Endogenous mouse Ig was blocked by incubating sections with donkey anti-mouse Fab fragments (Jackson Immunoresearch) diluted 1:50 in Biocare Medical Rodent Block M for 60 minutes. Sections were washed three times in TBS-T, then incubated with anti-mouse Fc for 15 minutes, before a further three washes in TBS-T. Non-specific antibody binding was inhibited by incubation with Biocare Medical Background Sniper with 1% BSA and 20% donkey serum and 20% goat serum for 15 minutes. Primary antibody 4G4 was diluted 3:4 in the above buffer and incubated on sections for 1 hr at room temperature. Sections were washed three times in TBS-T, then goat anti-mouse Perkin Elmer Opal HRP Polymer diluted 1:500 in TBST was applied and then bound to OPAL 570 for 60 minutes, before a further three washes in TBS-T. For NeuN, antigen retrieval was performed in Dako Epitope Retrieval Solution (pH 6.0) buffer at 125°C for 5 min. Endogenous mouse Ig was blocked by incubating sections with donkey anti-mouse Fab fragments (Jackson Immunoresearch) diluted 1:50 in Biocare Medical Rodent Block M for 60 minutes. Sniper/ 2% BSA block. Mouse anti-NeuN was diluted 1:500 in the above buffer and incubated on sections for 1 hr at room temperature. Sections were washed three times in TBS-T, then goat anti-mouse Perkin Elmer Opal HRP Polymer diluted 1:500 in TBST was applied and then bound to OPAL 520 for 60 minutes. Sections were washed three times in TBS-T, then incubated with DAPI (1:10,000) for 5 minutes, before a further three washes in TBS-T.

### RNA-seq and bioinformatics

RNA was extracted from infected mouse brains (as described above) using TRIzol (Invitrogen) or the Maxwell® RSC simplyRNA Tissue Kit (Promega), both as per manufacturer’s instructions. Viral RNA was extracted from virus stock culture supernatants using NucleoSpin RNA Virus kit (Machery Nagel) as per manufacturer’s instructions. RNA from the mouse brains (extracted using TRIzol) and virus stock was DNase I treated and cleaned using the RNeasy MinElute Cleanup kit (QIAGEN), as per manufacturers’ instructions. RNA concentration and quality was measured using TapeStation D1K TapeScreen assay (Agilent). RNA was prepared for sequencing using the Illumina Stranded Total RNA Prep Ligation with Ribo-Zero Plus (dual 10bp indexes) kit (Illumina Inc) and sequenced using the Illumina Nextseq 2000 platform. Per-base sequence quality for >90% bases was above Q30 for all samples. Raw sequencing reads were assessed using FastQC (v0.11.8)^35^ and MultiQC (v1.7)^36^ and trimmed using Cutadapt (v2.3)^37^ to remove adapter sequences and low-quality bases. Trimmed reads were aligned to the GRCm39 vM26 reference genome for mouse datasets, using STAR aligner (v2.7.1a)^38^, with more than 95% of reads mapping to protein coding regions. Viral genomes were *de novo* assembled from RNA-Seq libraries derived from virus-infected cell culture supernatant, using metaSPADes v3.15.5^39^ after QC and trimming as above. Viral reads in mouse samples were quantified by aligning reads to the JEV GII FU strain reference genome (Genbank AF217620.1)^40^ or to the appropriate (i.e. strain-specific) *de novo* assembled viral genome using Bowtie2 v2.2.9^41^, and counting ‘primary proper paired alignments’ with Samtools v1.10^42^. Gene expression was calculated using RSEM v1.3.1^43^. Differentially expressed genes (DEGs) were identified using DESeq2 v1.46.0 in R v4.4.1 as described above for Pseudo-bulk analysis, with sex included in the DESeq2 model. Data were analysed in R using the following R packages: dplyr (v1.4.4), ggpubr (v0.6.1), gplots (v3.2.0), Openxlsx (v4.2.8), PCAtools (v2.18.0), RColorBrewer (v1.1-3), tidyr (v1.3.1), tidyverse (v2.0.0).

### Gene set enrichment analysis

Using DESeq2 as described above, a log_2_ fold-change ranked gene list was generated for each group and a Gene Set Enrichment Analysis was performed using GSEA (v4.4.0)^44^ with 1000 permutations, the ‘no_collapse’ setting, and a set range of 5-1500 genes per set. The MSigDB v2023_1_Mm gene set collection, comprising of 15,579 *Mus musculus* gene sets, was obtained from the molecular signatures database (MSigDB)^45^. For comparing GSEA results between groups, gene sets that were significantly enriched in one group but not another were given a NES of 0 for the non-significant group.

### Pathway Analysis

Pathway analysis was performed using Ingenuity Pathway Analysis (IPA) v65367011 (QIAGEN) with default settings for ‘expression batch analysis’ using the log_2_ fold-change of the DEGs identified in DESeq2, as described above. Significantly activated/inhibited Canonical Pathways were identified using -log(p) > 1.3 and then ranking by z-score.

Significantly activated/inhibited Upstream Regulators were identified using p < 0.05 and ranking by z-score. All annotations lacking a z-score were ignored.

### Protein-protein interaction network analysis

A list of DEGs uniquely upregulated in the day 21 symptomatic group were analysed using the STRING database v12. Clustering was performed using the Markov Cluster Algorithm in the STRING web-app. Centrality metrics were calculated, and plotting performed, using Cytoscape v3.10 and the CytoHubba plugin.

### Apoe genotyping of a clinical Acute Encephalitis Syndrome (AES) cohort

Theoretical trypsin cleavage peptides were computed using the PeptideMass tool at https://web.expasy.org with the human APOE-ε3 (Protein ID: P02649) protein sequence from https://www.ncbi.nlm.nih.gov to create a panel of APOE SNP-associated tryptic peptides: CLAVYQAGAR (ε2-specific; labelled apoe2), LGADMEDVCGR (shared ε2/ε3; labelled apoe2_3), LGADMEDVR (ε4-specific; labelled apoe4), and LAVYQAGAR (shared ε3/ε4; labelled apoe3_4).

A set of 148 publicly available LC-MS/MS raw data files generated from cerebrospinal fluid (CSF) samples of patients with acute encephalitis syndrome (AES)^46^ were downloaded from the PRIDE repository (Accession: PXD034789) and re-analysed. Raw data were processed in Proteome Discoverer (v3.2.0.450; Thermo Fisher Scientific) using TMTpro 16-plex quantification. Peptide-spectrum matches were searched against the reviewed UniProt human reference proteome supplemented with APOE isotype sequences. Trypsin was specified as the protease with up to two missed cleavages. Methionine oxidation was included as a dynamic modification; cysteine carbamidomethylation and TMTpro modification of lysine residues and peptide N-termini were treated as static modifications. Searches were restricted to precursor charge states 2-4, precursor mass range 350-5000 Da, and peptide length 7-30 amino acids. Mass tolerances were 10 ppm for precursor ions and 0.02 Da for fragment ions. False discovery rate was controlled at 1% at the PSM, peptide, and protein levels. Reporter abundances were derived automatically using signal-to-noise (S/N) values when available, otherwise intensities were used.

To correct for loading error, peptide abundances were normalised against the reference channel within each TMT batch with the formula ‘Log_2__sample_channel - Log_2__reference_channel’. TMT batch effects were removed using ComBat from the SVA package. The effect of TMT batch-correction was examined using ANOVA to test for differences in mean peptide abundance between TMT batches with and without batch-correction. The effect of total APOE protein abundance on SNP-associated peptide abundance was removed using a linear model with the formula ‘Log_2__APOE_peptide ∼ Log_2__APOE_protein’. Using residuals from the linear model, samples were clustered using a 3-dimensional Gaussian mixture model with the mclust package. Feature selection (i.e. which and how many SNP-associated peptides to include in the model) and the optimal number of clusters (within in a range of 2 to 5) was determined using Bayesian Information Criterion. To create an expression profile for each cluster, SNP-associated peptide abundances were converted to global z-scores (i.e. across all clusters) and then averaged within each cluster. APOE genotypes were then assigned to each cluster based on mean z-score profiles. Where zygosity could not be confidently resolved, genotypes were combined (i.e. ε2/ε3 and ε2/ε2 were collapsed into ‘ε2/ε3 or ε2/ε2’, and ε4/ε3 and ε4/ε4 were collapsed into ‘ε4/ε3 or ε4/ε4’). APOE genotype frequencies of the general Laotian population^47^ were summed across ethnicities, collapsed into the same genotype groups as the AES cohort, and compared to the AES cohort using Fisher’s exact test in GraphPad Prism v10.

Categorical and binary clinical features recorded in the AES metadata (including CSF colour, survival, aetiology, seizures) were tested for disproportionate representation across APOE genotype groups using Fisher’s exact tests in GraphPad Prism. For numeric clinical features (including Days of illness, Glasgow Coma Scale on admission, Blood WCC, Blood CRP, CSF white cell count, CSF opening pressure, CSF neutrophil count, CSF lymphocyte count, CSF red cell count, and duration of admission), pairwise genotype comparisons were screened by effect size using Cohen’s *d,* and the three features with the greatest *d* were tested for equal variance using an F-test, and for differences between means with a two-sample *t*-test. p-values were adjusted for multiple testing using the Benjamini-Hochberg procedure. All analysis was done using R v4.4.2 unless noted otherwise.

### Software and packages

R version 4.4.2 with: Seurat v5.0, SeuratObject, FastReseg, DESeq2, clustree, scCustomize, ComplexHeatmap, pheatmap, EnhancedVolcano, Matrix, data.table, spatstat.explore, tidyverse (dplyr, ggplot2, readr, tidyr), cowplot, ggpubr, patchwork, openxlsx, RColorBrewer, gprofiler2.

### Statistics

The t-test was used when the difference in variances was <4-fold, otherwise the non-parametric Kolmogorov-Smirnov exact test or Mann-Whitney test was used (GraphPad Prism 8). Kaplan-Meier statistics were determined by log-rank (Mantel-Cox) test. The pertinent statistics for the bioinformatic data are described in the relevant methods sections or figure legends.

Weight-loss trajectory was modelled using a 3-parameter exponential growth curve *a + b^c×T^*, where *a* is the asymptote, *b* is the scale factor, *c* the growth rate and *T* the time. The parameters *a*, *b*, and *c*, were compared between groups using standard least squares for *a*, Negative LogNormal Maximum Likelihood for *b*, and LogNormal Maximum Likelihood for *c*. These analyses were conducted in *JMP Pro (v 18.2.2 SAS Institute, Cary NC)* using the *Fit Curve* platform.

The fit of the curves was assessed visually and via AIC, BIC, and R^2^ (0.94). Multi-modal curve fit was performed based on the distribution of the percentage consistent peak to trough weight loss. The Normal 2 Mixture and Normal 3 Mixture distributions are defined by two or three location μi, scale σi, and proportion πi parameter sets. These flexible distributions are capable of fitting bimodal or multi-modal data. Confidence intervals for normal mixture distribution parameter estimates use Wald-based calculations. AICc calculated in the JMP Distribution platform gives a measure of the goodness of fit of an estimated statistical model that can be used to compare two or more models. AICc is a modification of the AIC adjusted for small samples. AICc can be computed only when the number of data points is at least two greater than the number of parameters. AICc weights give normalised AICc values that sum to one. The AICc weight can be interpreted as the probability that a particular model is the true model given that one of the fitted models is the truth. Therefore, the model with the AICc weight closest to one is the best fit. The AICc weights are calculated using only non-missing AICc values: AICcWeight = exp[-0.5(AICc-min(AICc))] / sum(exp[-0.5(AICc-min(AICc))]) where min(AICc) is the smallest AICc value among the fitted models. These analyses were conducted in *JMP Pro (v 18.2.2 SAS Institute, Cary NC)* using the *Distribution* platform.

## RESULTS

### Spatial transcriptomics reveals immune activation and neuronal stress in lethal JEV infection, with near restoration of homeostasis in recovered mice

We have previously reported C57BL/6J mouse models of JEV and MVEV infection and disease that recapitulate key features of human disease^11^. Following subcutaneous inoculation, a subset of mice developed lethal encephalitis, with clear viral protein staining and histopathological lesions in the brain, indicating blood brain barrier crossing and direct infection of neurons. Other mice experienced substantial body weight loss but ultimately survived and recovered. Herein, we sought to determine the transcriptional signatures associated with lethal and non-lethal JEV by using single cell spatial transcriptomics of formalin fixed paraffin embedded (FFPE) mouse brains^11^. We opted for the CosMx platform with the 950-plex Mouse Neuroscience Panel spiked with probes for viral RNA (Supplementary Table 1), which was the largest multiplex panel available for mouse brain analysis at the time of the experiment. Samples included mouse brains from two uninfected controls, four with lethal infection (two JEV_Nayakama_, one JEV_FU_, and one MVEV_TC123130_), and one that recovered from symptomatic JEV_FU_ infection and harvested on day 30^11^. Unsupervised clustering at the optimised resolution (see Methods and Supplementary Figure 1) revealed 22 distinct cell clusters (Figure 1A-C). The neuronal clusters were largely segregated according to brain region (Figure 1B and Supplementary Figure 2). In mice that succumbed to lethal infection, JEV and MVEV were consistently detected by IHC in neurons of the cortex, and often also in the thalamus or caudate putamen^11^ (Figure 1D). Neurons infected by JEV were positive for both Rbfox3 transcripts by CosMx and NeuN protein by IHC (Supplementary Figure 3A,B), with Rbfox3/NeuN widely recognised as a marker of mature neurons^48^. Detection of viral RNA by CosMx (Figure 1E) was consistent with IHC (Figure 1D). Neurons were the main target of infection, particularly cortical glutamatergic neurons, caudate putamen neurons, and thalamic neurons (Figure 1F and Supplementary Figure 3C). While IHC of human brain sections is sparse, infection in cortical and thalamic neurons^49,50^ parallels with our mouse models.

**Figure 1.**
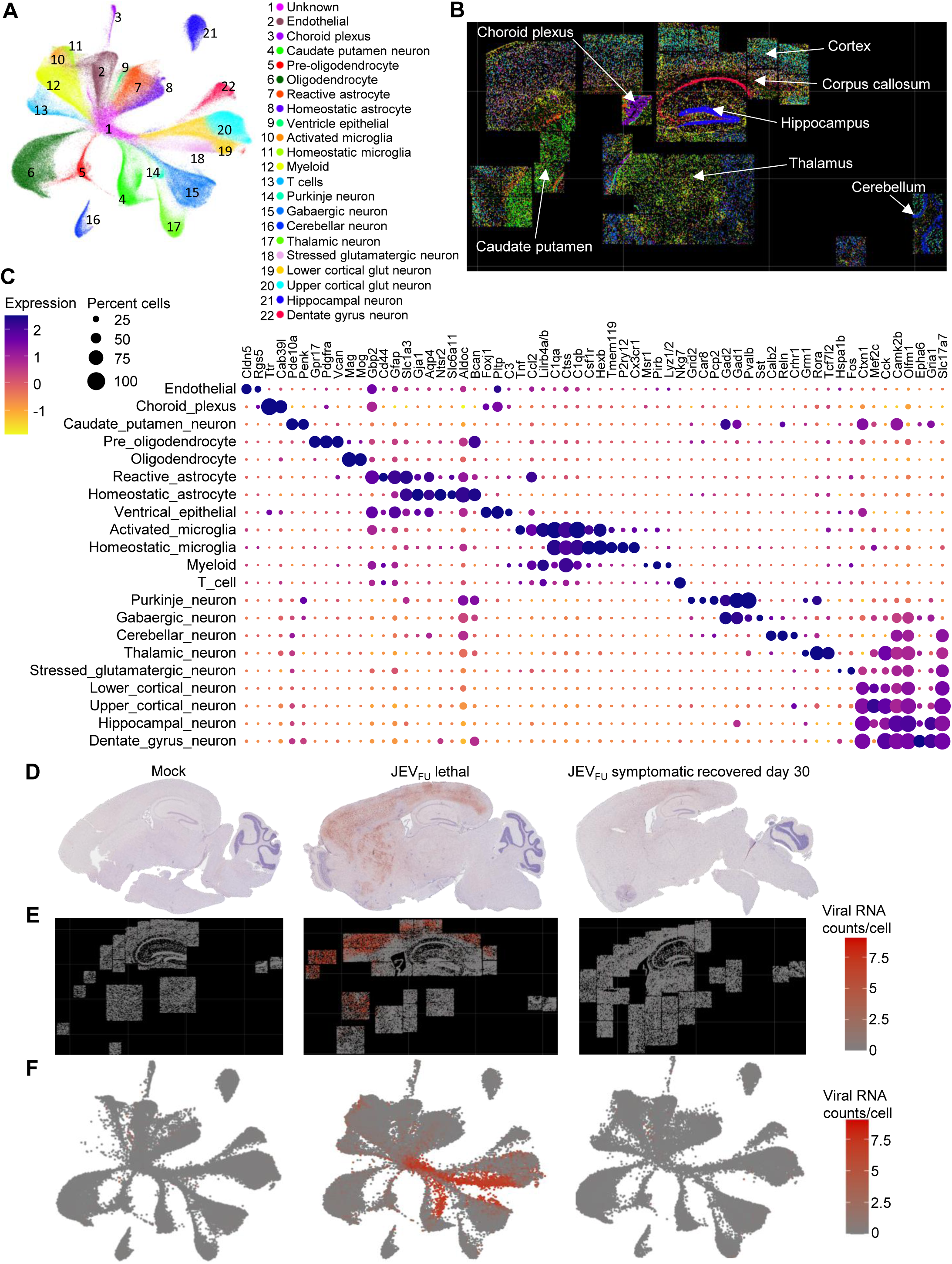
Single cell spatial transcriptomics of JEV infected mouse brains. A) UMAP plot displaying the integration of 232,164 cells from seven mouse brains (2 uninfected, 2 JEV_Nakayama_ lethal, 1 JEV_FU_ lethal, 1 MVEV_TC123130_ lethal, and 1 JEV_FU_ symptomatic recovered day 30 post infection). One unknown cluster and 21 cell-type clusters were identified using Leiden clustering, including neurons, astrocytes, oligodendrocytes, microglia, endothelial, and immune infiltrates (myeloid and T cells). B) Spatial distribution of cell-type clusters in a representative lethal mouse brain (MVEV_TC123130_), with major brain regions annotated. C) Dot plot of scaled expression levels (blue = high, yellow = low, circle size = percentage of cells with at least one count of the corresponding transcript) for distinguishing markers of the 21 cell types. Markers include both predicted cluster-specific markers and established canonical markers for each cell type. D) IHC staining (brown) of mouse brains (left = uninfected, middle = JEV_FU_ lethal, right = JEV_FU_ symptomatic recovered day 30 post infection) for *Orthoflavivirus* NS1 using the 4G4 monoclonal antibody. E) Per cell quantity and spatial distribution of JEV RNA reads for the corresponding mouse brains in ‘D’. F) UMAP plot showing per cell quantity of JEV RNA reads for the corresponding mouse brains in ‘D’ and ‘E’. Cell-type clusters are defined in ‘A’. Red shading corresponds to JEV RNA levels, where a more intense red indicates more counts.

Principal component analysis (PCA) revealed that the four lethal mouse brains (two JEV_Nakayama_, one JEV_FU_, and one MVEV_TC123130_) clustered closely together across all cell types (Supplementary Figure 4). These samples were therefore grouped and collectively analysed as lethal infection in subsequent analyses. The symptomatic recovered brain (day 30) clustered between the lethal-infected and mock brains for most cell types (Supplementary Figure 4).

JEV/MVEV infection triggered a substantial influx of myeloid cells, which became the dominant population (∼17% of total cells) in the brains of mice that succumbed to lethal infection (Figure 2A). In symptomatic mice that survived, myeloid cells accounted for ∼6% at day 30 post infection, compared with <1% in uninfected mice. T cells were virtually absent (<0.1%) in uninfected brains but represented ∼4% of total cells in both lethal and recovered brains (Figure 2A). In lethal cases, homeostatic microglia and astrocytes shifted into activated or reactive states, whereas in recovered mice these populations had largely returned to homeostasis by day 30 (Figure 2A). Infected cortical neurons in lethal brains exhibited a stressed phenotype driven by *Fos* and *Hspa1b* expression^51,52^, which was not observed in the recovered mice on day 30 post infection (Figure 2A). Together, these findings indicate that mice succumbing to lethal infection experience pronounced immune activation and neuron stress in the brain, whereas surviving mice show substantial restoration of homeostasis by day 30, despite persistent immune infiltrates.

**Figure 2.**
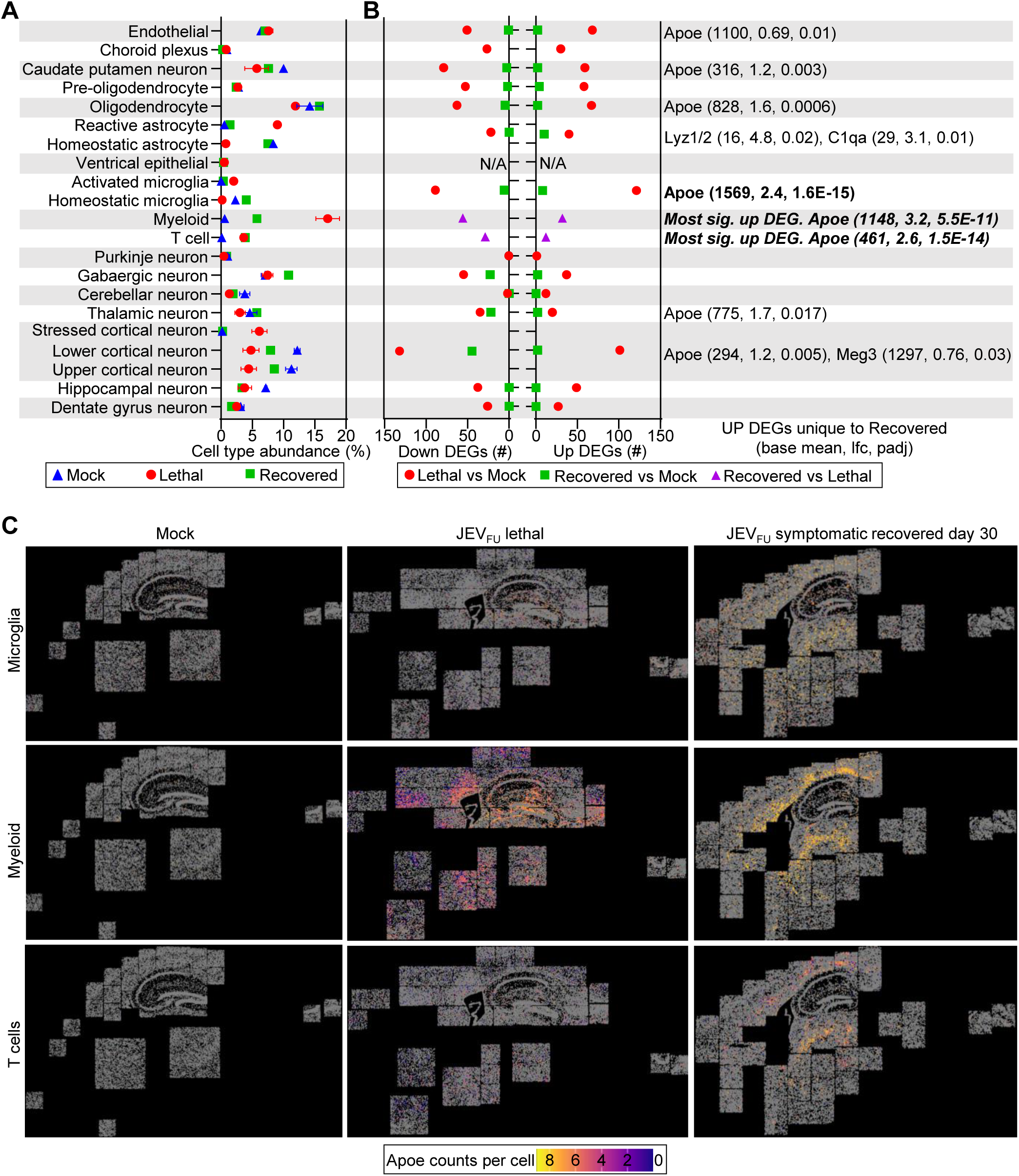
Pseudobulk differential expression analysis identifies *Apoe* as significantly upregulated in JEV symptomatic recovered mouse brains. A) Abundance of the 21 cell types as a percentage of total cells per brain section (blue triangles = mock, average ± SEM of n=2, red circles = lethal orthoflavivirus infection, average ± SEM of n=4, green squares = JEV_FU_ symptomatic recovered day 30 post infection, n=1). B) Pseudobulk differential expression analysis for each cell type comparing lethal (red circles) or symptomatic recovered (green squares) mice with mock controls. For myeloid and T cells, which had insufficient cell numbers in mock brains, symptomatic recovered mice were compared with lethal mice (purple triangles). The number of upregulated and downregulated DEGs are shown for each comparison and cell type. For analysis, homeostatic and activated microglia were combined into a single microglia group, homeostatic and reactive astrocytes into one astrocyte group, and cortical neuron sub-clusters were consolidated. N/A indicates insufficient cell numbers for robust statistical analysis. DEGs significantly upregulated in symptomatic recovered versus mock, but not in lethal versus mock, are listed by cell type, with base mean, log_2_ fold change, and adjusted p-value shown in brackets. For myeloid and T cells, the most significantly upregulated DEG is shown in italics. Raw data is in Supplementary Tables 2, 3 and 4. C) For cell types in which *Apoe* was the most significantly upregulated unique DEG in symptomatic recovered mice (microglia, myeloid, and T cells), per-cell *Apoe* transcript counts are shown for representative mouse brains (left = uninfected, middle = JEV_FU_ lethal; right = JEV_FU_ symptomatic recovered at day 30 post-infection). Blue shading corresponds to a low *Apoe* count per cell, pink-orange shading corresponds with a medium *Apoe* count per cell, and yellow shading corresponds to a high *Apoe* count per cell.

### Apoe was significantly upregulated in brain immune cells of symptomatic mice at day 30 post-infection, but not in mice with lethal infection

We performed pseudobulk differential expression analysis for each cell cluster, comparing lethal versus mock and recovered versus mock. For infiltrating immune populations (T cells and myeloid cells), which were absent in mock brains, we instead compared recovered versus lethal. To enable comparisons across conditions, homeostatic and activated microglia were combined into a single microglia group, homeostatic and reactive astrocytes were combined into one astrocyte group, and cortical neuron sub-clusters were consolidated. Across all cell types, lethal mouse brains exhibited more differentially expressed genes (DEGs) than symptomatic recovered brains at day 30 (Figure 2B). Microglia and cortical neurons showed the largest transcriptional changes in lethal brains, reflecting a pronounced pro-inflammatory state (Figure 2B, Supplementary Figure 5, Supplementary Table 2). In recovered mice, the most prominent changes were downregulated genes in neuron clusters susceptible to JEV infection, consistent with a neuroprotective phenotype. For example, *Egr1* was the most significantly downregulated gene in cortical neurons (Supplementary Table 3), and its loss has been shown to reduce cell death during viral encephalitis^53^. The limited number of upregulated genes in recovered mice largely overlapped with those in lethal brains, suggesting residual low-level inflammation. However, some genes were uniquely upregulated in recovered brains at day 30, with *Apoe* dominating across multiple cell types (Figure 2B, Supplementary Table 3, Supplementary Figure 6). This upregulation was most pronounced in immune cells, including microglia and infiltrating myeloid and T cells (Figure 2B-C).

Overall, the most significant unique transcriptional signature in symptomatic recovered mice at day 30 post-infection was the marked upregulation of *Apoe* in microglia and infiltrating immune cells.

### Brain microglia and infiltrating immune cells adopt an anti-inflammatory and resolving phenotype on day 30 in recovered mice compared to lethal infection

Apolipoprotein E (APOE) is a multifunctional protein involved in lipid transport between cells throughout the body^54^. It also plays important immunomodulatory roles, including promoting macrophage polarisation toward an anti-inflammatory M2 phenotype^55,56^ and regulating antigen presentation and T-cell priming^56^. In the homeostatic brain, *Apoe* was expressed most highly in astrocytes^54^ (Supplementary Figure 7). However, in mice that survive and recover from JEV brain infection, *Apoe* was also highly expressed in immune cell populations. In the recovered brain, *Apoe* expression is higher in myeloid cells and microglia than in astrocytes (Supplementary Figure 7), and is significantly upregulated in T cells, myeloid cells, and microglia compared with lethal infection (Figure 3A-C, Supplementary Table 4). In recovered brains, *Ccl2* was the most significantly downregulated gene across T cells, myeloid cells, and microglia, alongside other pro-inflammatory genes such as *Tnf*, *Stat2,* and *Ifng* (Figure 3A-C, Supplementary Table 4). Ingenuity Pathway Analysis (IPA) Upstream Regulators (USRs) further demonstrated a marked anti-inflammatory profile in these immune cell populations, characterised by upregulation of anti-inflammatory IL10 and IL1RN signalling, and downregulation of pro-inflammatory IFNG, TNF, IL1A and IL1B signalling in recovered versus lethal brains (Figure 3D). The APOE USR was also significantly upregulated in T cells, myeloid cells, and microglia (Supplementary Table 5), indicating functional pathway-level relevance. We also identified cell type-specific USRs. Erythropoietin (EPO), which promotes Treg proliferation^57^, was significantly upregulated in T cells (Figure 3D), suggesting a more Treg-like phenotype. TIMP1 (tissue inhibitor of metalloproteinases 1), which supports myelopoiesis^58^, was significantly downregulated specifically in myeloid cells (Figure 3D), supporting the reduced abundance of myeloid cells in the recovered versus lethal brain. The significant USRs aligned with canonical pathway analyses, which showed neuroinflammatory signatures were downregulated across T cells, myeloid cells, and microglia in recovered versus lethal brains, while resolution pathways were upregulated (Supplementary Table 5). Additionally, T cell exhaustion signalling was specifically downregulated in T cells, while MHC class II antigen presentation was specifically upregulated in myeloid cells (Supplementary Table 5). Collectively, these findings indicate that T cells, myeloid cells, and microglia in the recovered brain adopt an anti-inflammatory, resolution-phase and tissue-repair phenotype, contrasting with the inflammatory profile observed in the lethal brain.

**Figure 3.**
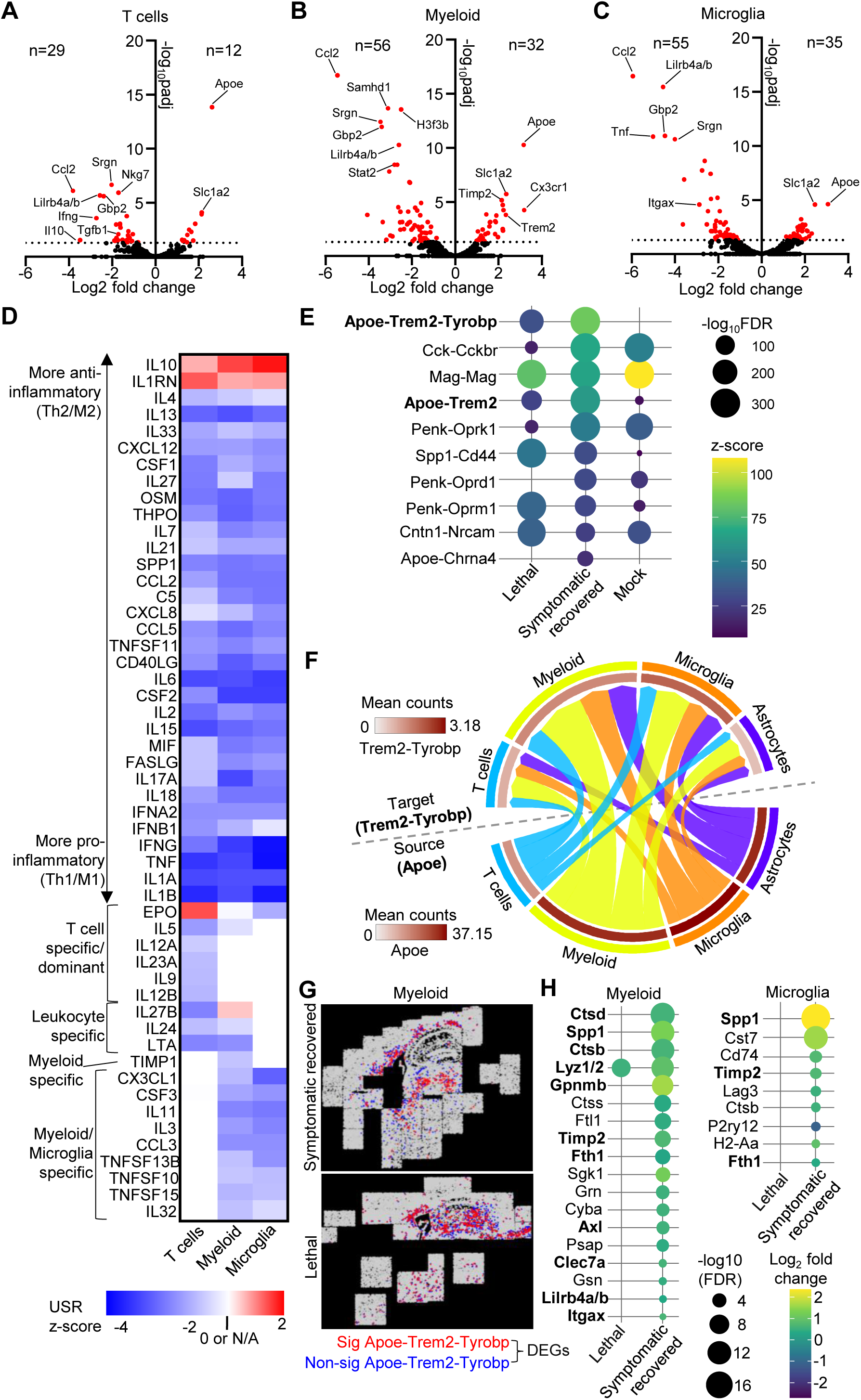
*Apoe-Trem2-Tyrobp* ligand-receptor interaction promotes phagocytic and anti-inflammatory states in myeloid cells and microglia during JEV recovery. A-C) Volcano plots showing log_2_ fold change and -log_10_ adjusted p-value for JEV symptomatic recovered day 30 versus lethal mouse brains for T cells (A), myeloid cells (B), and microglia (C). Dotted line represents threshold for significance (p<0.05), and significant genes are coloured red, with selected DEGs labelled. Number of DEGs are shown as n= on the left of the y-axis for downregulated DEGs and n= on the right of the y-axis for upregulated DEGs. Raw data is in Supplementary Table 4. D) The DEGs from ‘A-C’ (Supplementary Table 4) were analysed using Ingenuity Pathway Analysis (IPA) Upstream Regulator (USR) function, and the z-score for selected cytokine USRs (raw data is in Supplementary Table 5) are shown for T cells, myeloid cells, and microglia. Red indicates upregulation in JEV symptomatic recovered compared to lethal, blue indicates downregulation in JEV symptomatic recovered compared to lethal, and white indicates a z-score of 0 or not significant (N/A). E) Bubble plot showing the degree of spatial co-expression expressed as z-score (coloured scale) and significance (size of bubble, -log_10_ FDR) for the top 10 ligand-receptor interactions in the JEV_FU_ symptomatic recovered brain from spatialDM analysis. F) Cord diagrams showing ligand-receptor interactions in the recovered section involving *Apoe* and *Trem2-Tyrobp*. Outer ring shows the cell-type involved in each interaction. Inner ring is coloured by mean gene counts of ligand and receptor for each cell type. Cord width shows the relative number of significant interactions for each L-R/cell-type pair. The cell-types were chosen for plotting because they were the most highly represented source and target cell types among significant Apoe-Trem2-Tyrobp L-R interactions. G) Spatial distribution of ‘Apoe-receiver’ myeloid cells with significant (p<0.05) SpatialDM and *Apoe-Trem2-Tyrobp* receptor Moran’s I > 1 (significant, red), and ‘Apoe-non-receiver’ cells (not-significant, blue), for a JEV symptomatic recovered (top) and lethal (bottom) section. The same plots for microglia are shown in Supplementary Figure 9. H) Bubble plots showing differentially expressed genes between ‘Apoe-receiever’ versus ‘Apoe-non-receiver’ myeloid cells and microglia shown in Figure 3G and Supplementary Figure 9. Log_2_ fold change (coloured scale) and significance (size of bubble, - log_10_ FDR) for all DEGs in the JEV symptomatic recovered brain are shown, and all data is available in Supplementary Tables 6-7. Bolded genes are markers of ‘disease associated microglia’^60–62^.

### *Apoe-Trem2-Tyrobp* signalling promotes phagocytic and immunoregulatory states in myeloid cells and microglia during JEV recovery

To gain insight into cell-to-cell communication in JEV infected mouse brains, we performed ligand-receptor interaction analysis on the single cell spatial transcriptomics data. Of the 119 ligand-receptor pairs in the CosMx panel, 68 had significant global spatial co-expression in at least one condition (lethal, recovered, or mock) (Supplementary Table 6). In both the mock and lethal brains, the most significant receptor-ligand interaction was *Mag-Mag* (Figure 3E), which reflects oligodendrocyte myelin-axon maintenance signalling^59^. However, in the brains of JEV symptomatic mice at 30 days post-infection, the most significant ligand-receptor interaction was *Apoe-Trem2-Tyrobp* (Figure 3E). The *Apoe-Trem2* interaction alone, without the adaptor molecule *Tyrobp*, was also among the most highly significant ligand-receptor interactions observed in symptomatic recovering mice (Figure 3E). *Apoe-Trem2-Tyrobp* interactions were also significantly enriched in the JEV lethal brain, albeit at much lower magnitudes and significance (Figure 3E). In contrast, *Apoe-Trem2* was only marginally significant in uninfected mice, while *Apoe-Trem2-Tyrobp* did not reach statistical significance (Figure 3E). Most of the *Apoe-Trem2-Tyrobp* ligand-receptor interactions were between myeloid and microglial cells (Figure 3F), and this was also true in the lethal brain (Supplementary Figure 8C), but not the mock brain (Supplementary Figure 8B). Together, these data identify *Apoe-Trem2-Tyrobp* signalling between myeloid cells and microglia as a dominant cell-cell communication axis in the recovering JEV brain, while its lower-level presence during lethal infection suggests that this pathway may be engaged early but is amplified during resolution.

To define the transcriptional profile of cells receiving *Apoe* ligand via the *Trem2-Tyrobp* receptor complex, we compared receiver microglia and myeloid cells that were predicted to participate or not participate in this ligand-receptor interaction (Figure 3G). Participating and non-participating cells did not segregate into distinct cell type clusters (Supplementary Figure 9), indicating that engagement of the *Apoe-Trem2-Tyrobp* axis reflects a cell state transition rather than a completely distinct cell type. Differential expression analysis (Supplementary Table 7) revealed enrichment of genes associated with phagocytosis and lysosomal degradation (Ctsd, Ctsb, Ctss, Lyz1/2), iron sequestration and oxidative/ferroptotic stress buffering (Ftl1, Fth1), and immunoregulatory or suppressive functions (Gpnmb, Spp1, Cst7) (Figure 3H). Strikingly, most of these genes (Spp1, Gpnmb, Timp2, Axl, Clec7a, Lilrb4a/b, and Itgax) are established transcriptional markers of Apoe-Trem2-dependent “disease-associated microglia (DAM)”, a program induced by Apoe-Trem2 signaling following phagocytosis of apoptotic neurons in neurodegenerative settings^60,61^. In addition, other enriched genes (Ctsd, Ctsb, Lyz1/2, Fth1) are upregulated in early-stage DAMs that arise independently of Apoe-Trem2 signalling prior to full activation of DAMs^61^. DAMs exhibit enhanced phagocytic and lysosomal capacity and have been implicated in the clearance of pathological substrates such as amyloid plaques^61,62^. Together, these findings indicate that *Apoe-Trem2-Tyrobp* signaling in our dataset induces a conserved damage-response transcriptional program, suggesting that during viral encephalitis, not only microglia but also infiltrating myeloid cells can adopt a DAM-like state in an *Apoe-Trem2-Tyrobp* dependent manner during resolution and recovery.

### Establishing a model to interrogate early stages of resolving Japanese encephalitis

Neuroinflammation resolution is particularly difficult to study in humans due to limited access to brain tissue at critical timepoints spanning onset, progression, and recovery. Using mouse brain samples, we identified a distinct resolution-phase phenotype in myeloid cells and microglia from recovered mice at 30 days post-infection, defined by enhanced phagocytic and anti-inflammatory transcriptional signatures mediated through the *Apoe-Trem2-Tyrobp* signalling axis. However, by day 30 post-infection, inflammation resolution is well underway, meaning the critical early triggers and signalling events of the resolution process may be missed. We therefore sought to establish a mouse model of Japanese encephalitis resolution that could be monitored in real time, enabling sampling at the earliest stages of this process. The JEV_FU_ isolate, a genotype 2 strain isolated from human serum during a 1995 outbreak in Australia^63^, had a 20-30% mortality rate in C57BL/6J mice and has not been serially passaged in suckling mouse brains^11^. We thus chose JEV_FU_ for our study, because our other JEV isolates, JEV_Nakayama_ and JEV_NSW/22_, have been serially passaged in mouse brains or are not neuroinvasive in C57BL/6J mice, respectively^11^. The JEV_FU_ strain also recapitulates key human disease outcomes (lethal, symptomatic, and asymptomatic), which was not evident for JEV_Nakayama_ or JEV_NSW/22_^11^.

We subcutaneously infected 6-week-old C57BL/6J mice with 5×10^3^ CCID_50_ JEV_FU_, and monitored body weight, disease scores, and viremia. All mice developed a viremia peaking on days 2-3 and becoming largely cleared by days 4-5, with no differences in viremia between mice with subsequent different clinical outcomes (Figure 4A). Approximately 30% of mice developed weight loss and clinical scores that met humane euthanasia criteria (Figure 4B), based on a previously reported scorecard that predicts the final day of likely survival^11^. These cases are herein designated as having ‘lethal’ outcomes. Lethal outcomes occurred between 8- and 15-days post-infection (Figure 4B) and was associated with a rapid decline in body weight (Figure 4C, lethal). Mice that survived infection had widely variable changes in body weight, with some losing ∼19% body weight before recovering, and others not losing any weight (Figure 4C). For the following experiments, we defined symptomatic recovered mice as those that exhibited a consistent peak-to-trough weight loss of at least 5%, which was the maximum weight fluctuation observed in uninfected mice, followed by subsequent weight gain. Mice that did not show a consistent peak-to-trough weight loss of 5% or more were classified as asymptomatic. Once mice begin to recover from body weight loss during the typical window for lethal outcomes (approximately days 8-17), they are unlikely to succumb to infection^11^. Brain samples were collected for histological analyses (left hemisphere) and RNA-seq (right hemisphere) under seven scenarios: (i) uninfected mice (n=7), (ii) mice with lethal outcomes (n=17), (iii) symptomatic mice one day post-peak weight loss (n=6), (iv) symptomatic mice on day 21 (n=6), (v) symptomatic mice on day 30 (n=8), (vi) asymptomatic mice on day 21 (n=10), and (vii) asymptomatic mice on day 30 (n=4) (Figure 4C). The ‘day post-peak weight loss’ group was chosen to capture the earliest signatures of inflammation resolution and recovery. Earlier signatures of inflammation resolution and recovery cannot be determined, as this is the earliest timepoint at which we can confirm that mice will not succumb to JEV infection. The day 21 timepoint represents an intermediate stage between the earliest possible timepoint (day post-peak weight loss) and the symptomatic recovered day 30 timepoint which was analysed by single cell spatial transcriptomics (Figures 1-3).

**Figure 4.**
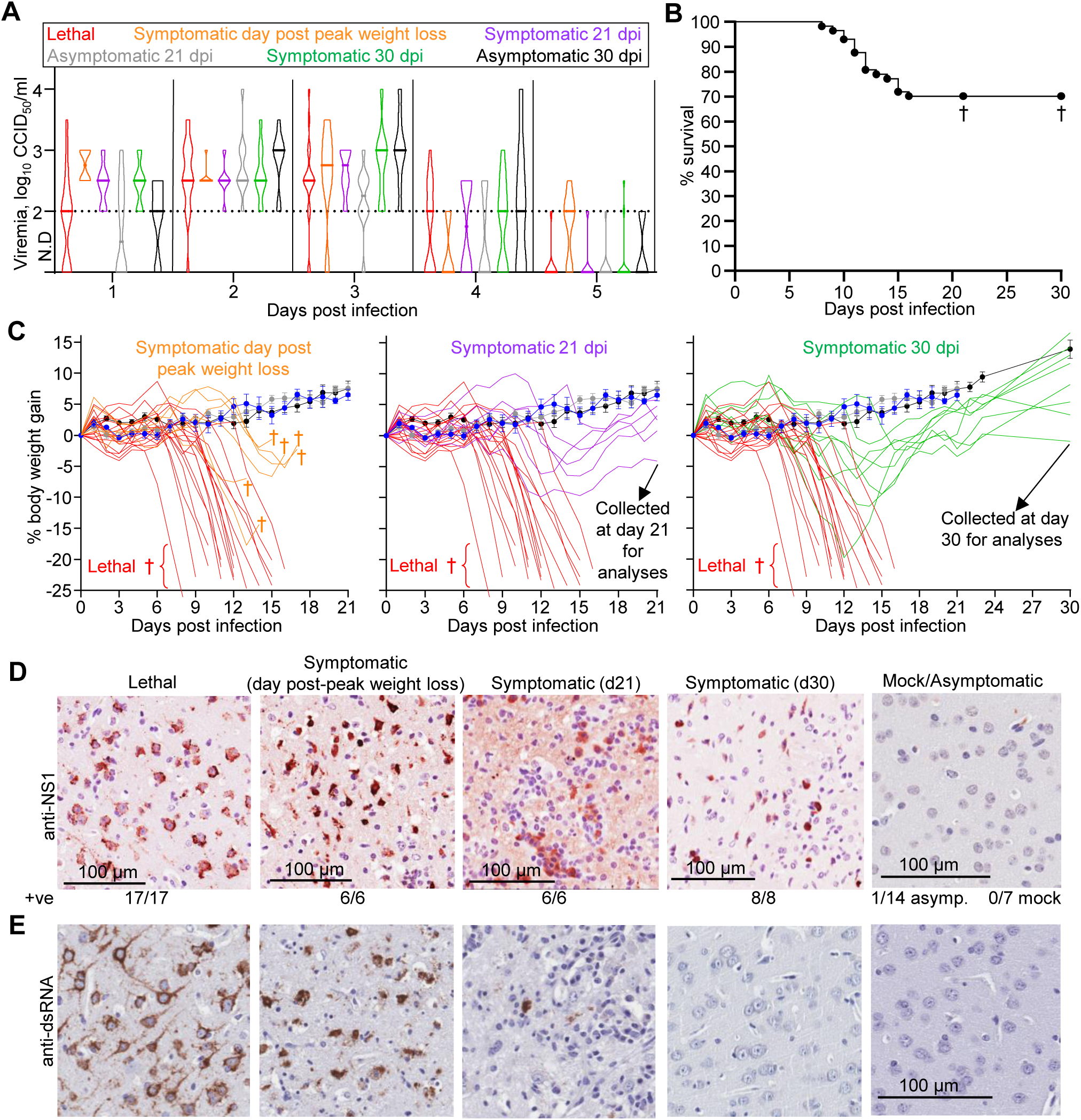
Establishing a mouse model of resolving Japanese Encephalitis. 57 female ≈6-week-old C57BL/6J mice were infected s.c. with 5 × 10^3^ CCID_50_ of JEV_FU_. Mock mice (n=4) were injected s.c. with diluent (RPMI + 2% FBS) and collected on day 21 (RNA-seq batch 2, see Figure 5D), or were not injected and collected (n=3) for comparison with lethal and day 30 post-infection (see RNA-seq batch 1, Figure 5D). A) Violin plots showing daily serum viremia (CCID_50_/ml) over 5 days for lethal (red, n = 17), symptomatic day post-peak weight loss (orange, n=6), symptomatic day 21 (purple, n=6), asymptomatic day 21 (grey, n=10), symptomatic day 30 (green, n=8), and asymptomatic day 30 (black, n=10). The horizontal line within the violin plot represents the median. All mice recorded a detectable viremia for at least one timepoint. Limit of detection is 2 log_10_CCID_50_/ml of serum (dotted line). B) Kaplan–Meier plot showing percent survival (n=57). Mice euthanised at pre-determined time points without reaching the humane endpoint disease score (see methods) were classified as survivors. † denotes the day 21 and day 30 timepoints. C) Percent body weight change of mice after infection with JEV_FU_ at 5 × 10^3^ CCID_50_ compared to each mouse’s weight on day zero. Lethal or symptomatic mice (those that exhibited a consistent peak-to-trough weight loss of at least 5%) are shown as individual data, while mock and asymptomatic mice (those that did not that did not show a consistent peak-to-trough weight loss of 5% or more) are shows as mean ± SEM. Group colour and mouse number corresponds to panel ‘A’, except n=3 mock mice for comparison at day 30 post infection were not weighed. Mice were euthanised at one day after peak weight loss (left panel, orange), day 21 (middle panel, purple, grey, and blue), and day 30 (right panel, green, and black), or when they reached the score requiring humane euthanasia (lethal, red). D) IHC staining (brown) of mouse brains for *Orthoflavivirus* NS1 using the 4G4 monoclonal antibody. IHC images are representative of C57BL/6J mice with lethal (n = 17), symptomatic day post-peak weight loss (n=6), symptomatic day 21 (n=6), symptomatic day 30 (green, n=8), asymptomatic day 21 (grey, n=10), asymptomatic day 30 (black, n=4), and mock (n=7). Numbers of mice with clear positive staining (out of the total per group) are indicated below the images. E) Images of the corresponding representative mouse brains from panel ‘D’ with IHC (brown) staining for double-stranded RNA (dsRNA) using an anti-dsRNA monoclonal antibody.

To determine the level of JEV infection in the mouse brains, paraffin sections were stained with anti-non-structural protein 1 (NS1) or anti-double-stranded RNA (dsRNA) monoclonal antibody. All lethal and symptomatic recovered mice exhibited JEV neuroinvasion, as evidenced by detection of cells positive for JEV NS1 (Figure 4D). In symptomatic recovered mice, JEV NS1 staining typically co-localised with increased Iba1 staining (Supplementary Figure 10), indicating the localised presence of microglia and/or infiltrating myeloid cells. Gradual clearance of virus from the brain was observed, with dsRNA (a marker of active viral replication) cleared more rapidly than NS1 (Figure 4E). Of the 14 asymptomatic mice, only one showed detectable JEV NS1 staining, suggesting that the asymptomatic phenotype occurred because the virus did not cross the blood brain barrier or infect neurons. Overall, we have established a mouse model in which weight loss serves as a reliable real-time marker of Japanese encephalitis, enabling accurate identification of the earliest timepoints of inflammation resolution for sampling and analysis.

### Early resolution-phase RNA-Seq signatures reveal enhanced phagocytosis, antigen cross-presentation, anti-inflammatory polarisation, and cholesterol metabolism

We performed bulk RNA-seq on mouse brains collected at lethal endpoints, one day post peak weight loss, day 21 (symptomatic and asymptomatic), and day 30 (symptomatic and asymptomatic), and compared these to uninfected controls. Mice that succumbed to lethal JEV infection had viral reads in the brain ranging from 0.45% to 18% of total reads (Figure 5A). Three non-synonymous mutations were identified in JEV reads in two mice that died from infection; one mouse had G153W mutation in envelope and K40E in NS1, and another mouse had L31F in NS3 (Supplementary Table 8). G153W may impact the N154 glycosylation site, which has been linked to pathogenicity^64^. Otherwise, there were no consistent JEV mutations associated with lethal brain infection. On average, brains from mice at lethal endpoints had significantly higher levels of viral RNA than those collected one day post peak weight loss, which in turn had significantly higher levels than symptomatic mice at day 21 (Figure 5A). At both days 21 and 30, symptomatic mice had significantly higher viral RNA levels than asymptomatic and uninfected mice, respectively (Figure 5A). These findings are consistent with IHC results (Figure 4D) and confirm JEV neuroinvasion followed by progressive viral clearance in symptomatic mice undergoing recovery. PCA further demonstrated clear segregation between groups, with symptomatic mice clustering closer to lethal mice than the asymptomatic mice, which clustered more closely with uninfected controls (Figure 5B). This indicates substantial transcriptional differences between symptomatic and asymptomatic mice sampled at matching timepoints. Consistent with this, there were only 2 DEGs between asymptomatic and mock mice at both day 21 and day 30, compared with 1294 DEGs for symptomatic versus mock at day 21, and 232 DEGs at day 30 (Supplementary Table 9). Earlier during resolution, at one day post peak weight loss, there were 4337 DEGs, and at lethal endpoints there were 8820 DEGs (Supplementary Table 9). Overall, these results further indicate that transcriptional responses to JEV neuroinvasion occur in symptomatic mice, but not in asymptomatic mice.

**Figure 5.**
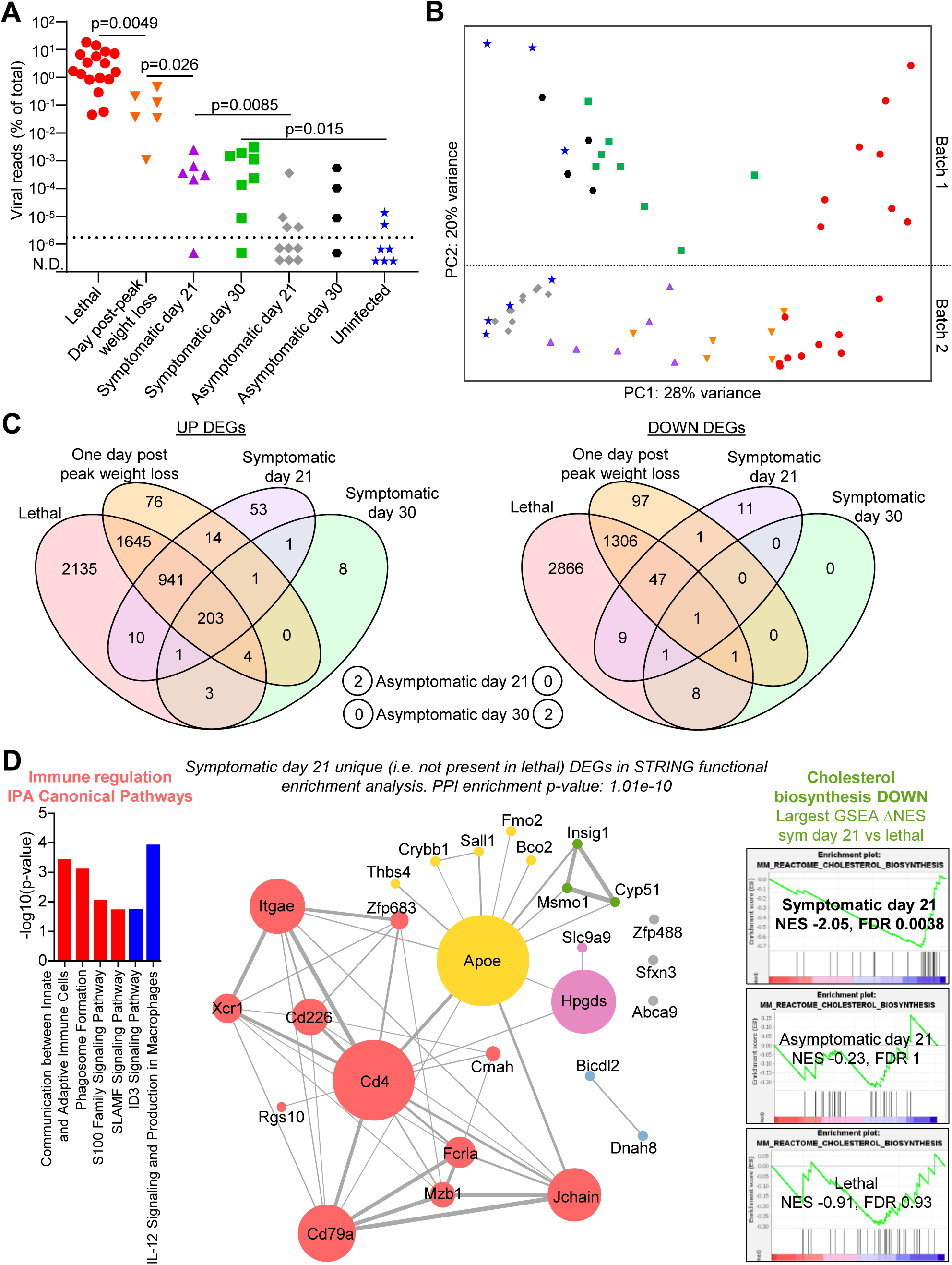
Bulk RNA-Seq analyses of C57BL/6J mice infected with JEV_FU_. C57BL/6J mouse brains (right hemisphere) were subjected to bulk RNA-seq for mock mice (blue stars, n=7), mice with lethal outcomes (red circles, n=17), symptomatic mice one day post-peak weight loss (orange downward triangles, n=6), symptomatic mice on day 21 (purple upward triangles, n=6), symptomatic mice on day 30 (green squares, n=8), asymptomatic mice on day 21 (grey diamonds, n=10), and asymptomatic mice on day 30 (black hexagons, n=4). A) JEV RNA reads as a percent of total reads. Statistics by Kolmogorov-Smirnov exact test. B) Principal Components Analysis (PCA) based on normalised variance-stabilised transformed (vst) counts of all genes (excluding zero-variance genes) from bulk RNA-seq data. Samples above the dotted line were sequenced in batch 1, and below the dotted line in batch 2. Comparisons for DEG analyses were only performed within respective batches. C) Venn diagrams of upregulated (left) and downregulated (right) DEGs (adjusted p-value <0.05) for each condition versus mock. The lethal DEGs are a combination of DEGs in batch 1 and batch 2 to increase stringency for identifying unique DEGs in non-lethal conditions. D) Network analysis using the STRING database. Input was the 81 DEGs up- or down-regulated in symptomatic day 21 but not in lethal. STRING removed genes not in the database (unannotated or uncharacterised genes, and Ig genes), resulting in 27 genes for analysis. This network had a protein-protein interaction (PPI) enrichment p-value of 1.01e-10, indicating this network has significantly more interactions than would be expected for a random set of proteins. Colour of node indicates the functional cluster based on unsupervised MCL clustering in STRING (red = immune regulation, green = cholesterol biosynthesis, magenta = prostaglandins/endosomal ion transport, blue = motor protein cargo transport), yellow = Apoe and others. Node size represents log_2_ betweenness scores calculated using Cytoscape. Line thickness is the combined interaction score from the STRING database. IPA Canonical Pathway analysis of the 81 DEGs is shown on the left, with the -log10 p-value shown for all 6 significantly up- or downregulated pathways with z-scores (red = positive z-score, blue = negative z-score. Raw data in Supplementary Table 13). Gene set enrichment analysis (GSEA) plots are shown on the right for the MM_REACTOME_CHOLESTEROL_BIOSYNTHESIS geneset, which had the largest difference in normalised enrichment score (NES) between symptomatic recovered day 21 compared to lethal. NES and FDR are shown for lethal and day 21 symptomatic and asymptomatic.

Most of the up- and downregulated DEGs in symptomatic mice at one day post peak weight loss, day 21, and day 30 were also differentially expressed in mice with lethal outcomes (Figure 5C), indicating persistent encephalitis-associated signatures (e.g. *Ccl5*, *Oasl1*, *Isg15*, *Stat1*, *Irf9*, *Gfap*, see Supplementary Table 9 for full list) that gradually diminish over time. To specifically identify resolution-associated pathways, we focused on DEGs that were uniquely up- or downregulated in symptomatic recovering mice but not differentially expressed in lethal mice (Figure 5C). These unique DEGs likely represent inflammation resolution signatures that were not activated in mice with lethal encephalitis.

On day 30, there were only 10 uniquely upregulated DEGs (Supplementary Table 10) which were mostly related to astrocyte-mediated synapse homeostasis (Supplementary Figure 11A). Riluzole is an FDA-approved drug for the treatment of amyotrophic lateral sclerosis (ALS). Its pharmacological properties include: (1) inhibition of glutamate release, (2) inactivation of voltage-dependent sodium channels, and (3) interference with intracellular signalling events following activation of excitatory amino acid receptors. It has also demonstrated neuroprotective effects in various *in vivo* models of neuronal injury involving excitotoxicity^65–67^. Given that these mechanisms mirror several uniquely upregulated pathways identified in the brains of symptomatic mice at day 30 post infection (Supplementary Figure 11A), we evaluated whether Riluzole could improve JEV outcomes. When administered via drinking water starting one day prior to infection, Riluzole treatment led to increased viremia (suggesting potentially systemic immunosuppressive effects), but resulted in a significant reduction in weight loss among non-lethal cases (Supplementary Figure 11B-E). In contrast, when dosing commenced on day 6 post infection (after viremia had resolved but before lethal brain infection) there was no significant difference in viremia between groups, and no significant effect on weight loss or mortality (Supplementary Figure 11F-I). Although there were modest indications of protective effects leading to increased survival, these did not consistently reach statistical significance. Overall, Riluzole does not appear to be a viable therapeutic candidate for JEV, and astrocyte-mediated synaptic homeostasis may represent a late phase resolution process associated more with neuronal repair than with inflammation resolution and survival.

*Apoe* showed a log_2_ fold change of 1.1 in symptomatic mice at day 30 versus mock (compared with 0.05 in lethal (batch 1) and 0.21 in asymptomatic mice versus mock), although this did not reach statistical significance. However, *Apoe* expression was significantly higher in symptomatic day 30 mice compared to lethal mice (Supplementary Table 9), supporting the CosMx findings (Figures 2 and 3). In addition, Gene Set Enrichment Analysis (GSEA) using the full Molecular Signatures Database (MSigDB), which uses all genes ranked by fold change rather than DEGs^33,68,69^, identified the Apolipoprotein Binding gene set as significantly enriched at day 30 in symptomatic mice, but not in lethal mice (Supplementary Table 11-12), further supporting the CosMx results.

*Apoe* was significantly upregulated in symptomatic mice on day 21 (lfc 0.89, fdr 0.0006), but not at one day post peak weight loss (lfc 0.13, fdr 0.77) (Supplementary Table 9). Since day 21 was the earliest timepoint at which we detected *Apoe* upregulation, which was identified by our single cell spatial transcriptomics experiment as the most significant unique marker of recovery (Figure 2), we analysed the DEGs in symptomatic mice at day 21 that were not differentially expressed in lethal mice (i.e. ‘unique’) using the STRING database. These genes had a highly significant protein-protein interaction enrichment p-value, indicating that the proteins that these genes encode are at least partially biologically connected. Markov Clustering identified distinct gene clusters related to cholesterol biosynthesis and immune regulation (Figure 5D). Cholesterol biosynthesis genes *Msmo1*, *Cyp51* and *Insig1* were significantly downregulated in symptomatic mice brains at day 21 (Supplementary Table 10), and the Reactome Cholesterol Biosynthesis gene set was also significantly downregulated (Figure 5D). Unique DEGs at this timepoint also included *Cd4*, *Zfp683* (Hobit), *Itgae* (CD103), and *Cd226* which are markers of T cells including tissue resident memory T cells and Tregs^70–72^, *Xcr1* which is a marker of cross-presenting dendritic cells^73^, *Cd79a*, *Jchain*, and *Mzb1* which are markers of plasma B cells^74–77^, and *Rgs10* which is a key anti-inflammatory protein with important roles in M2 macrophage polarisation^78^. These findings are consistent with IPA Canonical Pathway analysis, which indicated significant upregulation of phagocytosis and antigen presentation pathways, alongside downregulation of pro-inflammatory signalling (Figure 5D). A similar signature was observed in mice one day after peak weight loss (Supplementary Figure 12, Supplementary Tables 9-13), although *Apoe* was not significantly upregulated at this time point (Supplementary Table 9). However, *Apoe* showed a modest log_2_ fold change of 0.126 (equivalent to approximately a 10% increase) which may be obscured in bulk RNA-seq due to dilution by cell populations that do not alter *Apoe* expression during JE resolution (i.e. non-microglia, non-myeloid, and non-T cells). Notably, for the unique DEGs at day 21 in symptomatic mice, *Apoe* displayed the highest betweenness centrality (Figure 5D, circle size), indicating it exerts the greatest influence among the unique DEGs in coordinating interactions between genes involved in immune regulation and cholesterol biosynthesis during JEV resolution and recovery. Cholesterol is synthesised locally in the brain, primarily by astrocytes, oligodendrocytes, and microglia, and ApoE serves as the key homeostatic regulator that binds to cholesterol and transports it to other brain cells to support remodelling and repair^79^. ApoE has also been shown to induce anti-inflammatory phenotypes in immune cells, either downstream of cholesterol signalling^80–82^, or via other mechanisms independent of lipid metabolism^55,83–85^. Overall, our integrated single-cell spatial transcriptomics and bulk RNA-seq analyses of mouse brains during JEV resolution revealed a defining molecular signature marked by myeloid and microglia ApoE engaging Trem2 to drive Tyrobp-dependent signalling and acquisition of a phagocytic, anti-inflammatory phenotype.

### *ApoE^-/-^* mice experienced enhanced JEV disease and were unable to recover due to dysregulated immune responses

To functionally test the role of ApoE in JE resolution, we infected *ApoE^-/-^* mice with JEV and examined clinical outcomes and brain inflammatory responses. There was no overt difference in viremia between wild-type C57BL/6J and *ApoE^-/-^* mice (Figure 6A). The survival rate of *ApoE^-/-^*mice infected with JEV was 58%, compared with 83% in wild-type C57BL/6J mice (Figure 6B), suggesting a trend to higher lethality although this difference was not statistically significant in the Kaplan-Meier survival analysis. However, *ApoE^-/-^*mice with lethal outcomes exhibited significantly faster weight loss compared to C57BL/6J mice with lethal outcomes (Figure 6C-D), suggesting more severe encephalitis. Rate of weight loss in C57BL/6J mice did not correlate with viral levels in the brain (Supplementary Figure 13A), indicating the rate of weight loss is not entirely dependent on viral load. A multi-modal curve fit of the percentage consistent peak to trough weight loss revealed the expected segregation of JEV-infected C57BL/6J mice into three groups: asymptomatic (minimal weight loss, no evidence of brain infection), symptomatic recovered (weight loss followed by weight gain, evidence of brain infection), and lethal (weight loss, disease score for humane euthanasia reached) (Figure 6E). In contrast, JEV-infected *ApoE^-/-^* mice lacked a symptomatic-recovered group, and instead all animals either remained asymptomatic or progressed to lethal infection (Figure 6D-E). Peak weight loss was significantly higher in non-lethal C57BL/6J mice compared to non-lethal *ApoE^-/-^* mice (Figure 6E), further underscoring the absence of a subgroup of mice that experiences and then clears brain infection in the *ApoE^-/-^* cohort. These findings suggest that in the absence of ApoE, once JEV gains access to and infects the brain, the virus is not efficiently cleared, and the ensuing encephalitis fails to resolve. This highlights a critical role for ApoE in orchestrating a non-lethal inflammatory response that both enables effective viral clearance and allows the subsequent resolution of inflammation.

We performed bulk RNA-seq on mouse brains collected during lethal encephalitis and at day 21 post-infection to assess transcriptional changes associated with ApoE deficiency. Viral read counts did not differ significantly between C57BL/6J and *ApoE^-/-^* mice with lethal outcomes (Figure 6F), suggesting the faster weight loss in *ApoE^-/-^*mice was not due to increased viral load. Six C57BL/6J mice that survived had viral reads above background levels (Figure 6F), and these mice had weight loss indicative of a symptomatic infection (Figure 6D-E). There were no non-lethal *ApoE^-/-^* mice with viral reads above background levels, and viral reads were significantly lower than for non-lethal C57BL/6J mice (Figure 6F). The non-lethal C57BL/6J mice with significant weight loss and viral reads also clearly segregated by PCA, while all *ApoE^-/-^* mice clustered either with the asymptomatic/mock mice or the lethal mice (Figure 6G). These data further support the idea that, without ApoE, JEV that gains access to the brain is not effectively cleared and the associated encephalitis does not resolve.

To pinpoint the ApoE-dependent signalling defects underlying the failure to resolve inflammation and the resulting increase in disease severity, we compared the brain transcriptomes of lethal *Apoe^-/-^* and C57BL/6J mice (Figure 6H). Of the 255 DEGs, 244 were downregulated in *Apoe^-/-^* mice (Figure 6I). *Apoe* was by far the most significantly downregulated gene (Figure 6I), validating the knockout mice. Most other significantly downregulated DEGs in *Apoe^-/-^* mice were associated with immune system functions (Supplementary Table 14). The observed downregulation of immune system functions in *Apoe^-/-^* mice was not attributable to reduced leukocyte infiltration, as H&E analysis showed a modestly increased overall abundance of leukocyte infiltrates compared with C57BL/6J controls, although this difference did not reach statistical significance (Supplementary Figure 13B), and in addition Iba1 IHC (myeloid and microglia) was not significantly different (Supplementary Figure 13C). Notably, *Il10ra*, the receptor for the anti-inflammatory cytokine IL-10 (whose signalling was upregulated in T cells, myeloid cells, and microglia during resolution in C57BL/6J mice (Figure 3B)), was among the most significantly downregulated genes in *Apoe^-/-^* mice (Figure 6I). *C5ar2*, a key mediator of pro-inflammatory Th1 contraction via IL-10 signalling^86,87^, was also markedly downregulated (Figure 6I). Together, the coordinated reduction of *Il10ra* and *C5ar2* suggests a diminished capacity to engage IL-10 mediated immunoregulatory pathways in the absence of ApoE, potentially impairing normal inflammatory resolution. MHC Class II genes (*H2-Aa*, *H2-Eb1*, and *Cd74*, among others) were significantly downregulated in *Apoe^-/-^* mice (Figure 6I), with phagocytosis and antigen presentation also a key resolution signature on day 21 (Figure 5D) which may be impaired in the absence of ApoE. Pathway analyses revealed an overall dysregulation of leukocyte activation, recruitment, antigen presentation, phagocytosis, and T-cell priming in *Apoe^-/-^* mice (Figure 6J and Supplementary Table 15).

Several genes related to lipid handling and metabolism (*Apoe*, *Apoc2*, *Apol7e*, *Trem2*, *Sgpl1*, *Lpcat2*, *Alox5ap*, *Lsr*, *Acox3*, *Pllp*) were significantly downregulated in *Apoe^-/-^*mice with lethal JEV (Supplementary Table 14). Apart from *Apoe*, none of the other genes were differentially expressed between asymptomatic *Apoe^-/-^*and asymptomatic C57BL/6J mice, indicating that disruptions in lipid-handling and metabolism in the brain arise specifically during encephalitis. Consistent with this, lipid handling and metabolism pathways were downregulated in IPA Canonical Pathways, Diseases and Functions, Gene Ontology, and Gene Set Enrichment Analysis (Supplementary Table 15). However, while lipid handling and metabolism pathways were clearly affected, their overall contribution to the transcriptomic changes was relatively minor compared with the more extensive dysregulation of immune responses.

Given the strong indication from transcriptomic analyses that *Apoe^-/-^* mice with lethal JEV infection exhibit a dysregulated immune response, we next quantified key immune cell populations in the brain using histology and IHC. However, shifts in the proportions of specific cell types are often more informative. An increased ratio of neutrophils (Ly6G⁺) to monocytes/macrophages/microglia (F4/80⁺) has been associated with pro-inflammatory states in multiple disease settings^33,88–91^. In *Apoe^-/-^* mice, we observed a modest increase in Ly6G^+^ neutrophils and a corresponding modest decrease in F4/80^+^ monocytes/macrophages/microglia (Figure 6K), though neither change reached statistical significance individually. However, this pattern was also observed in our RNA-seq results (Supplementary Table 14), where Ly6G transcripts were slightly upregulated (log_2_FC = 0.4, FDR = 0.88) and F4/80 (ADGRE1) transcripts were significantly downregulated (log_2_FC = - 1.06, FDR = 0.008). When we calculated the Ly6G^+^ / F4/80^+^ cell ratio for each mouse based on IHC quantification, *Apoe^-/-^* mice showed a significantly higher ratio compared to C57BL/6J controls (Figure 6L). Together, these findings support the interpretation that *Apoe* deficiency leads to a dysregulated immune response and skews the immune environment toward a more pro-inflammatory profile, with neutrophils contributing prominently to this shift.

**Figure 6.**
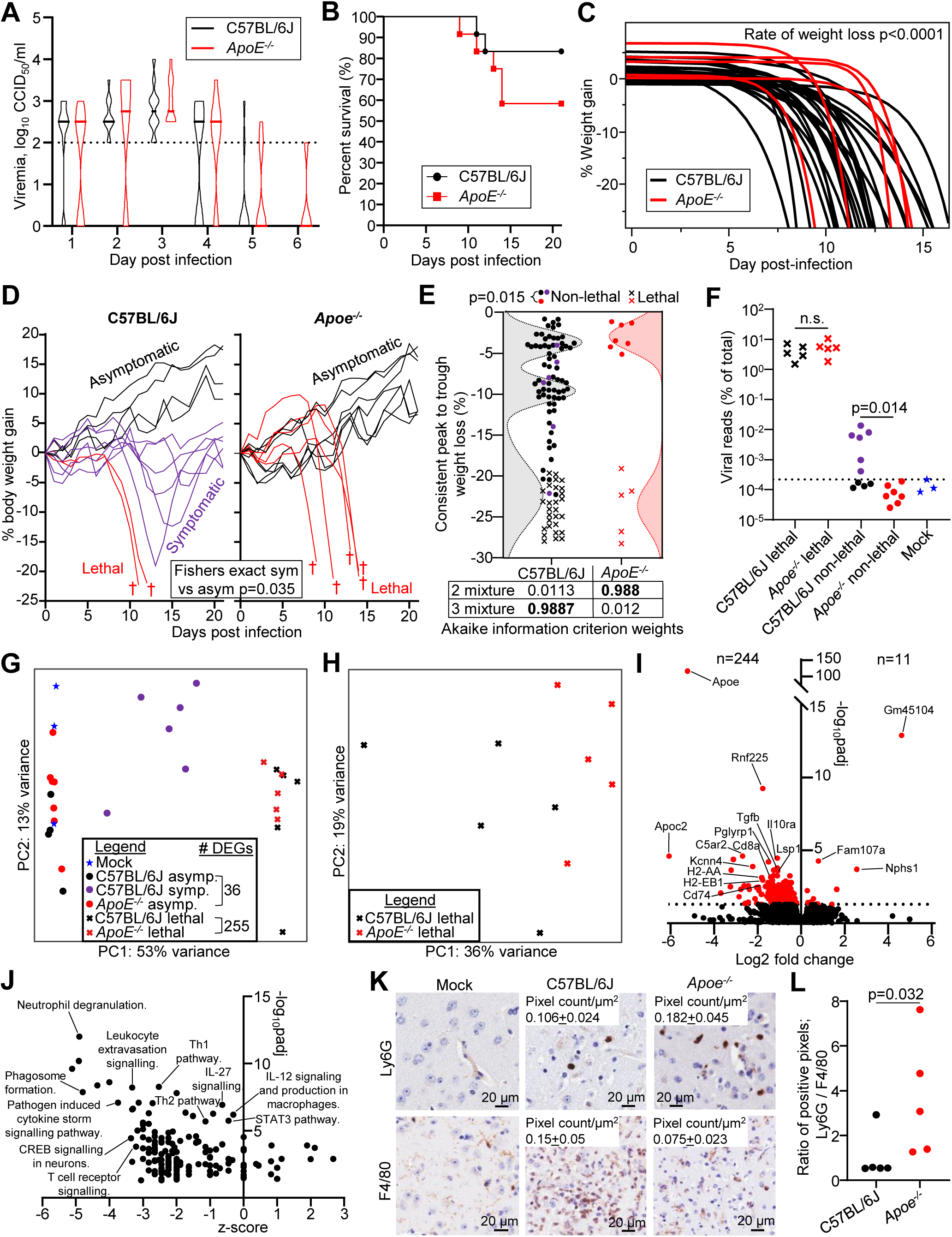
*ApoE^-/-^* mice exhibit more severe JEV disease and fail to recover from brain infection. 6-week-old C57BL/6J (n=6 males and n=6 females) and *ApoE^-/-^* (n=6 males and n=6 females) were infected s.c. with 5 × 10^3^ CCID_50_ of JEV_FU_. A) Violin plots showing daily serum viremia (CCID_50_/ml) over 5 days for C57BL/6J (black, n=12) and *Apoe^-/-^* (red, n=12) mice. The horizontal line within the violin plot represents the median. All mice recorded a detectable viremia for at least one timepoint. Limit of detection is 2 log_10_CCID_50_/ml of serum (dotted line). B) Kaplan-Meier plot showing percent survival for C57BL/6J (black, n=12) and *Apoe^-/-^* (red, n=12) mice. C) For each mouse (C57BL/6J lethal, black lines, n = 28 which includes n=2 from Figure 6D, n=17 from Figure 4C, and n=9 from Supplementary Figure 11D and 10H, and *ApoE^-/-^* lethal, red lines, n = 5), the weight-loss trajectory was modelled using a three-parameter exponential growth curve. Statistics is a t-test of growth rate for C57BL/6J versus *Apoe^-/-^* mice. D) Percent body weight change of individual C57BL/6J mice (n = 12) and *Apoe^-/-^* mice (n = 12) after infection with JEV_FU_ at 5 × 10^3^ CCID_50_ compared to each mouse’s weight on day zero. Two C57BL/6J and five *ApoE^-/-^* (red) reached the pre-determined disease score (see methods) and were euthanised (†). Six C57BL/6J mice had symptomatic brain infection (purple), while all non-lethal *Apoe^-/-^* were asymptomatic (black). Fisher’s exact test was used to compare asymptomatic and symptomatic mice across C57BL/6J and *Apoe^-/-^* mice. E) Consistent peak to trough weight loss percentage for each mouse (C57BL/6J, black, n = 99 which includes n=51 from Figure 4C, n=36 from Supplementary Figure 11D and 10H, and n=12 from Figure 6D, and *ApoE^-/-^* red, n = 12 from Figure 6D). Mice that survived (non-lethal) are indicated by filled circles, and mice that reached the endpoint for humane euthanasia (lethal) are indicated by crosses. Purple filled circles represents JEV-positive C57BL/6J symptomatic mice from panels ‘F’ and ‘G’. Statistics are by Kolmogorov-Smirnov exact test comparing all non-lethal mice for C57BL/6J versus *ApoE^-/-^*. The dotted curve (black = C57BL/6J, red = *ApoE^-/-^*) shows the multi-modal curve fit based on the distribution of the percentage consistent peak to trough weight loss. The best fitting curve is shown (i.e. Normal 3 Mixture distribution for C57BL/6J and Normal 2 Mixture distribution for *ApoE^-/-^*), as determined by the Akaike information criterion (AICc) weights. F) JEV RNA reads from bulk RNA-seq as a percent of total reads. Purple filled circles represents C57BL/6J non-lethal mice with viral reads above uninfected samples, denoted C57BL/6J symptomatic in panel ‘G’. Statistics by Kolmogorov-Smirnov exact test. G) PCA based on normalised variance-stabilised transformed (vst) counts of all genes (excluding zero-variance genes) from bulk RNA-seq data. The number of DEGs for asymptomatic *Apoe^-/-^* versus C57BL/6J was 36, and lethal *Apoe^-/-^* versus C57BL/6J was 255. H) PCA based on normalised variance-stabilised transformed (vst) counts from bulk RNA-seq data of the top 1000 most variable genes between lethal *Apoe^-/-^* and C57BL/6J. I) Volcano plots showing log_2_ fold change and -log_10_ adjusted p-value for lethal *Apoe^-/-^* versus C57BL/6J. Dotted line represents threshold for significance (p<0.05), and significant genes are coloured red, with selected DEGs labelled. Number of DEGs are shown as n= on the left of the y-axis for downregulated DEGs and n= on the right of the y-axis for upregulated DEGs. Raw data is in Supplementary Table 14. J) Volcano plots showing z-score and -log_10_ adjusted p-value for IPA Canonical Pathways generated from the DEGs for lethal *Apoe^-/-^* versus C57BL/6J. Selected pathways are labelled, and raw data is in Supplementary Table 15. K) IHC staining for neutrophil Ly6G and monocyte/macrophage/microglia F4/80. Images are representative of the n=3 mock and n=5 C57BL/6J and *Apoe^-/-^* lethal mouse brains from the RNA-seq panels ‘F’ and ‘G’. Medium and strong positive pixels were calculated using the Aperio Positive Pixel Count Algorithm v9, and the mean pixel count per µm and standard error are shown. L) The ratio of Ly6G^+^ / F4/80^+^ positive pixels (medium + strong) for each mouse. Statistics by Mann-Whitney test.

### APOE ε2 carriers have significantly reduced CSF neutrophils and hospitalisation duration

In humans, ApoE exists as three functionally distinct isoforms (APOE ε2 (Cys112/Cys158), ε3 (Arg112/Cys158), ε4 (Arg112/Arg158))^92^. In multiple neurological disease contexts, particularly Alzheimer’s disease, APOE ε2 is protective, ε3 is neutral, and ε4 is pathogenic^92^. Bharucha *et al*. performed a proteomic analysis of cerebrospinal fluid from 148 patients with Acute Encephalitis Syndrome (AES), including 60 patients with Japanese encephalitis (JE) and the remainder with viral, bacterial, or fungal aetiologies^46^. We reanalysed this dataset to infer the APOE isoform genotype of each patient. APOE SNP-associated tryptic peptides (Figure 7A, Supplementary Figure 14) were detected in all samples. Three-dimensional plotting of z-scores for peptide sequences that are distinct for APOE ε2, APOE ε4 and shared between APOE ε2/ε3 (Figure 7A) resolved three distinct clusters corresponding to APOE ε2 carriers (ε2/ε3 or ε2/ε2), APOE ε4 carriers (ε4/ε3 or ε4/ε4), and APOE ε3 homozygotes (ε3/ε3) (Figure 7B). The distribution of APOE genotypes did not differ significantly from that reported in the general Laotian population^47^ (Figure 7C), indicating that APOE isoform did not influence susceptibility to developing AES and supporting the accuracy of our allele assignments. APOE isoform frequencies were also not significantly different between patients who survived and those who died, although APOE ε2 carriers showed a modestly lower representation among fatal outcomes (Figure 7D). In contrast, APOE genotype was associated with disease recovery in AES survivors. APOE ε2 carriers had a significantly lower CSF neutrophil percentage compared to APOE ε3 homozygotes (Figure 7E). In addition, APOE ε2 carriers had a significantly shorter duration of hospitalisation compared with APOE ε4 carriers, while APOE ε3 homozygotes exhibited an intermediate duration (Figure 7F). Together, these findings suggest that the APOE ε2 isoform is associated with more rapid recovery from AES, perhaps driven by a blunted pathogenic neutrophil response, whereas APOE ε4 is associated with delayed recovery, consistent with the neuroprotective and neuropathogenic roles of these isoforms, respectively^92,93^.

**Figure 7.**
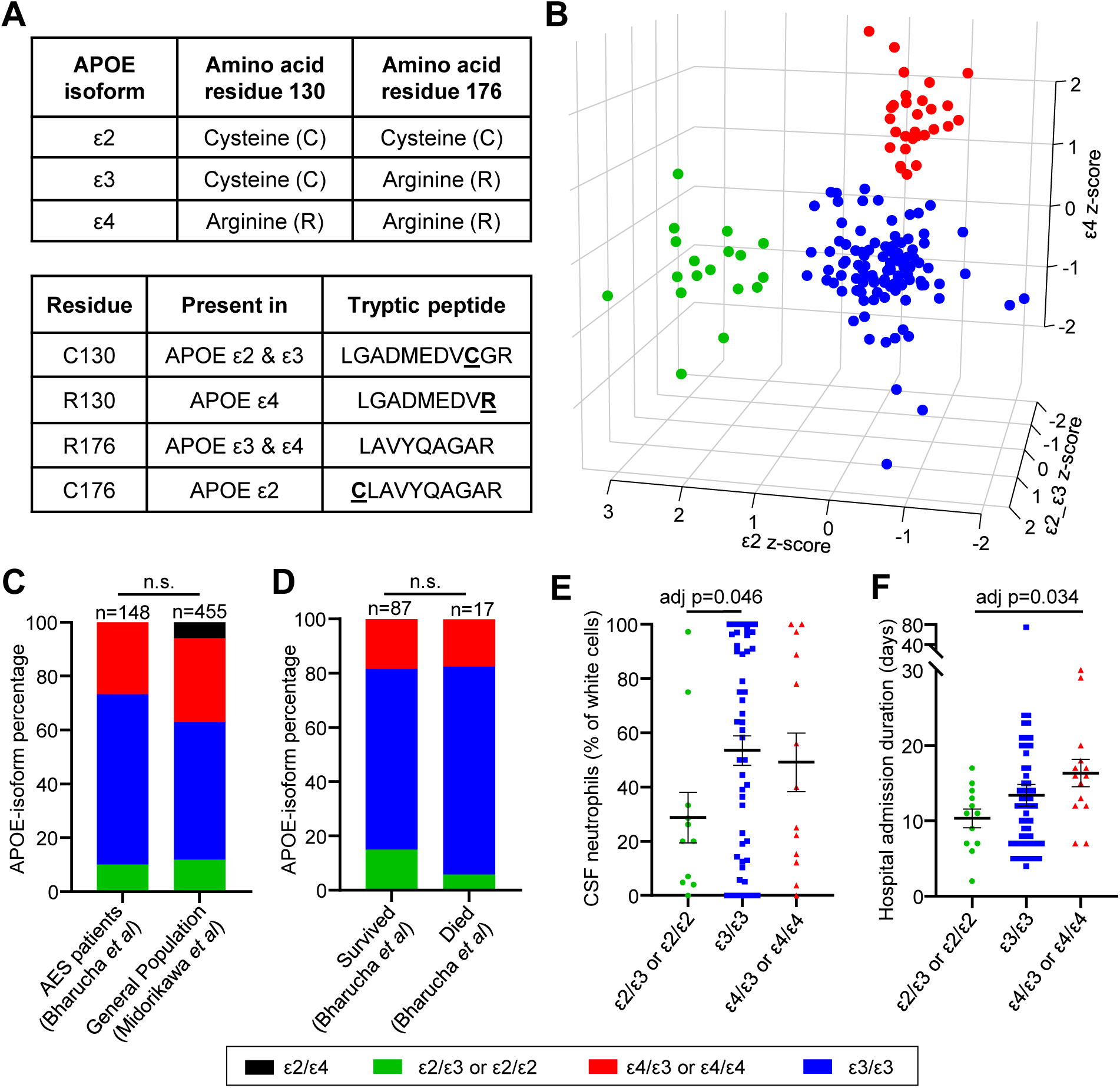
APOE-isoform assignment in AES patients reveals APOEε2 carriers have shorter hospital admission duration compared to APOE ε4 carriers. A) APOE isoforms (ε2, ε3, ε4) differ at amino acid positions 130 and 176. These isoforms are distinguishable in proteomics data by the abundance of SNP-associated tryptic peptides that are unique to APOE ε2 or APOE ε4, and those that are shared between APOE ε2 and ε3, and between APOE ε3 and ε4. B) Peptide abundances (z-scores) for peptides that are distinct for APOE ε2, APOE ε4 and shared between APOE ε2/ε3. Three distinct clusters corresponding to APOE ε2 carriers (ε2/ε3 or ε2/ε2, n=18, green), APOE4 carriers (ε4/ε3 or ε4/ε4, n=101, red), and APOE3 homozygotes (ε3/ε3, n=29, blue) were identified by Gaussian mixture modelling of normalised peptide abundances, and genotyped according to peptide z-scores. C) Percentage of each APOE-isoform in AES patients (n=148)^46^ and the Laotian general population (n=455)^47^. Proportions were not significantly different by Chi-square test or Fisher’s exact test. D) Percentage of each APOE-isoform in AES patients^46^ who survived (n=87) and those who died (n=17). Proportions were not significantly different by Chi-square test or Fisher’s exact test. E) CSF neutrophil percentage of total white cells. Samples with zero white cell count, or zero neutrophil and lymphocyte % were removed. Mean and standard error are shown. Statistics by t-test for APOE ε2-carriers versus APOE ε3 homozygotes. F) Duration of admission in hospital for AES patients who survived. ε2/ε3 or ε2/ε2 n=17, ε4/ε3 or ε4/ε4 n=14, ε3/ε3 n=54. Mean and standard error are shown. Statistics by t-test for APOE ε2-carriers versus APOE ε4-carriers.

## DISCUSSION

Among symptomatic Japanese encephalitis patients, approximately one third die, one third experience prolonged or permanent neurological sequelae, and one third make a full recovery^4^. Effective resolution of neuroinflammation and restoration of tissue homeostasis is critical for recovery after viral encephalitis. However, the immunological processes that govern inflammation resolution in *orthoflavivirus* encephalitis have not been systematically tracked or defined. In this study, we identify ApoE as a critical determinant of neuroinflammation resolution and recovery following JEV infection. By integrating a mouse model of Japanese encephalitis resolution, single-cell spatial transcriptomics, bulk RNA-seq, and functional genetics of human clinical data, we show that successful recovery is not defined solely by control of viral replication or passive decay of pro-inflammatory cytokines, but by the active engagement of a coordinated resolution program. Central to this program is the induction of ApoE in brain-resident and infiltrating immune cells, where it engages myeloid and microglial Trem2 to drive Tyrobp-dependent signalling and the transition from a pro-inflammatory to a pro-resolving state. Functional experiments demonstrate that ApoE is esential for this transition, where Apoe-deficient mice exhibit dysregulated immune responses, accelerated clinical deterioration, and a complete failure to recover following symptomatic infection. These findings indicate that ApoE is not merely a marker of recovery-associated cell states, but an essential component of the host response that enables restoration of tissue homeostasis.

ApoE is best known as a lipid shuttle that ferries cholesterol and other lipids between cells via members of the LDL-receptor family in the periphery and within the brain, thereby supporting membrane turnover, synaptic remodelling, and injury repair^54,79^. In symptomatic recovering mice, *Apoe* was upregulated alongside an overt downregulation of cholesterol biosynthesis signatures (Figure 5D), which may be triggered by JEV-induced IFN signalling^94^. This represents a pattern indicative of a shift from sterol production toward efflux and recycling^95^, which may also limit the accumulation of intracellular sterol species that can sustain inflammation, and simultaneously supplies redistributed cholesterol needed for tissue repair and recovery^79,95^. Consistent with ApoE’s role as the brain’s primary lipid-transport protein^96^, these coordinated changes suggest that ApoE-mediated lipid redistribution reduces the need for *de novo* cholesterol synthesis as part of the metabolic shift that accompanies successful resolution of Japanese encephalitis.

Beyond its role in lipid transport, ApoE serves as a key immunomodulator. Myeloid-derived ApoE, acting through members of the LDLR family and TREM2, promotes a shift from an M1 pro-inflammatory state toward an M2 resolving phenotype and suppresses the production of inflammatory cytokines^55,81,85,97,98^. This aligns closely with our observation that myeloid cells, T cells, and microglia in symptomatic recovering mice adopt an anti-inflammatory profile driven by IL10 signalling, whereas these same cell types display a pronounced pro-inflammatory state in lethal encephalitis, in which *Apoe* is not upregulated (Figures 2-3). Consistent with this immunoregulatory function, ApoE-deficient mice exhibit exaggerated systemic inflammatory responses (TNFα, IL-6, IL-12, and IFN-γ) following lipopolysaccharide (LPS) administration, an effect independent of hypercholesterolemia^83^. Moreover, ApoE restrains IL-12 production after stimulation with the TLR3 agonist poly I:C^83^. ApoE also binds activated C1q with high affinity, functioning as a checkpoint inhibitor of the pro-inflammatory classical complement cascade and attenuating unresolvable inflammation^84^. These findings indicate that ApoE limits Th1-polarising cytokine output, thereby suppressing excessive type I inflammatory responses *in vivo*^83^. This is consistent with our dataset that shows ApoE upregulation coinciding with the downregulation of the ‘IL-12 signalling and production in macrophages’ pathway specifically in recovering mice (Figure 5D). It also aligns with the reduced *Il10ra* and *C5ar2* and broad downregulation of pathways related to leukocyte activation and function at lethal endpoints (resolution failure) in *Apoe^-/-^*mice compared to wild-type mice (Figure 6I-J).

Myeloid-derived ApoE regulates antigen-presenting cell function by limiting membrane-cholesterol accumulation, thereby restraining MHC-II-dependent antigen presentation and reducing CD4⁺ T-cell priming, whereas ApoE deficiency enhances antigen presentation and T-cell activation in some settings^82^. ApoE also indirectly dampens T-cell activation by reducing surface densities of co-stimulatory and MHC-II molecules on antigen-presenting cells, leading to reduced IFN-γ responses^99^. Within the brain, ApoE interacts with microglial TREM2, promoting phagocytosis of ApoE-opsonised apoptotic neurons^96^. Our data shows that in symptomatic recovering mice, *Apoe* upregulation coincided with increased myeloid antigen MHC-II presentation and phagocytic signatures (Figure 3G, Figure 5D, Supplementary Table 5). In addition, *Apoe^-/-^* mice exhibited reduced expression of MHC-II genes and phagocytosis pathways (Figure 6I-J). Importantly, the transcriptional programs associated with *Apoe-Trem2-Tyrobp* signalling in our dataset closely resemble those of disease-associated microglia (DAM), suggesting that viral encephalitis and neurodegenerative disease may utilize a shared ApoE-Trem2 tissue-repair program. DAM arise in response to neuronal damage and are characterised by enhanced phagocytic and lysosomal activity alongside immunoregulatory functions, driven largely by APOE-TREM2 signalling, and are conserved in both humans and mice^60–62^. In line with this, *Apoe* upregulation in JEV recovering mice was associated with *Trem2-Tyrobp*-dependent induction of phagocytic, lysosomal, and DAM-associated genes, indicating that both microglia and infiltrating myeloid cells adopt a DAM-like state also during viral encephalitis resolution. Mechanistically, APOE-TREM2-TYROBP signalling promotes phagocytic activity via PI3K/AKT and PLCγ pathways^100,101^, while also exerting anti-inflammatory effects by modulating JNK and NF-κB signalling, thereby skewing cells toward a reparative phenotype and limiting pro-inflammatory cytokine production^102,103^. Consistent with our observation of increased ferritin light chain 1 (Ftl1) and ferritin heavy chain 1 (Fth1) in myeloid cells exhibiting strong *Apoe-Trem2-Tyrobp* interactions (Figure 3G), APOE signalling through the PI3K/AKT1 pathway can suppress ferroptosis by promoting ferritin (Fth1 and Ftl1) expression and limiting labile Fe^2+^ accumulation, while concurrently driving macrophage polarisation toward an anti-inflammatory M2 phenotype^85^. In neurodegenerative and injury models, these DAM pathways facilitate debris clearance, immune resolution, and tissue repair, whereas TREM2 deficiency or inhibition impairs phagocytosis and lysosomal function and exacerbates inflammation^104–106^. Consistent with these roles, *Apoe-Trem2-Tyrobp* signalling emerged as the most prominent receptor-ligand interaction between infiltrating myeloid cells and microglia during the symptomatic recovery phase of JE (Figure 3E-G). Although detectable at lower levels in lethal disease, its reduced magnitude suggests that while these resolution programs may be initiated during acute inflammation^107^, they are insufficiently amplified to drive DAM-like signatures (Figure 3G) needed for resolution and recovery. Together, our findings support a model in which ApoE upregulation during JE recovery promotes a DAM-like activation state in microglia and infiltrating myeloid cells via Trem2-Tyrobp signalling. This state enhances phagocytic clearance of cellular debris and immunogenic material, while promoting anti-inflammatory signalling. Through these coordinated functions, ApoE-driven DAM-like responses contribute to the termination of danger signalling, resolution of neuroinflammation, and restoration of brain homeostasis following encephalitis.

In a proteomics study of Japanese encephalitis patient cerebrospinal fluid, Yin *et al*. reported that individuals with less severe cognitive impairment had significantly higher APOE levels than those with more severe disease, suggesting that APOE may serve as a molecular marker of favourable JE prognosis^108^. This aligns with our findings that *Apoe* upregulation is a defining feature of successful inflammation resolution during recovery from Japanese encephalitis. Notably, the more severe (APOE^low^) patient cohort exhibited reduced proportions of immune infiltrates (neutrophils, monocytes, and lymphocytes), alongside increased classical complement activity, compared to the less severe (APOE^high^) cohort^108^. This is consistent with our observation that *Apoe^-/-^* mice show dysfunctional immune-infiltrate signatures (Figure 6J), and aligns with the known role of APOE in restraining classical complement signalling^84^. Similarly, in a proteomic study of CSF from patients with neuroinvasive West Nile virus encephalitis, APOE levels were significantly lower than in healthy controls, further supporting a pattern in which low APOE correlates with severe encephalitis^109^. In addition, another clinical study found that elevated peripheral pro-inflammatory cytokines (TNF-α and GM-CSF) were associated with increased lethality from Japanese encephalitis^110^. Together, these human and animal data reinforce a model in which ApoE abundance protects against severe neuroinflammatory outcomes, whereas ApoE deficiency or low expression correlates with heightened inflammation, complement overactivation, dysfunctional immune infiltration, and increased disease severity.

ApoE in humans has three functionally different protein isoforms (APOE ε2 (Cys112/Cys158), ε3 (Arg112/Cys158), ε4 (Arg112/Arg158))^111^. These amino acid variations alter the lipid transport and receptor binding capabilities of APOE. Peripherally, APOE ε4 facilitates efficient lipid clearance, APOE ε3 provides a balanced lipid exchange, whilst APOE ε2 clears lipids more slowly, leading to accumulation in the bloodstream^54,112,113^. In the brain, APOE ε2 promotes a neuroprotective environment, transporting lipids to neurons for membrane and synaptic repair. APOE ε3 functions neutrally, providing a balanced lipid exchange. APOE ε4 has reduced lipid delivery to neurons, weakens blood-brain barrier integrity, and promotes pro-inflammatory pathways, increasing the risk of neurodegenerative diseases (such as Alzheimer’s)^54,114–116^. Murine ApoE has only one allelic variant that functionally resembles human APOE ε3 and thus lacks the isoform diversity observed in humans^117^. Our reanalysis of publicly available proteomics data from human Acute Encephalitis Syndrome (AES) patients^46^ revealed significantly lower CSF neutrophil proportions and significantly shorter hospital admission duration in individuals carrying the APOE ε2 allele. Neutrophil proportions are a key marker of inflammation, and a high neutrophil-lymphocyte ratio is indicative of more severe encephalitis and poor prognosis^118^. Our findings indicate that APOE isoforms differentially modulate the magnitude of hyperinflammation and the trajectory of recovery during AES in humans. These findings are consistent with established isoform-specific effects of APOE on neuroinflammation, whereby APOE ε2 and APOE ε4 are associated with anti-inflammatory and pro-inflammatory glial signatures, respectively, in a TREM2-dependent manner^119^. This suggests that APOE isoforms may represent genetic risk factors for JE and AES inflammation severity and recovery. The association between APOE genotype and outcome in an aetiologically diverse acute encephalitis syndrome cohort raises the possibility that ApoE-dependent resolution pathways represent a conserved host response to neuroinflammation rather than a mechanism unique to JEV.

In summary, our data suggest that ApoE functions as an important driver of the transition from pathogenic to resolving neuroinflammation. ApoE upregulation in microglia and infiltrating myeloid cells is strongly associated with a shift toward phagocytic and anti-inflammatory immune states via engagement of *ApoE-Trem2-Tyrobp* signalling during JEV recovery. In contrast, ApoE deficiency is associated with dysregulated immune responses, leading to failed resolution and increased disease severity in mice. Human data from infectious AES patients further suggest that APOE isoform influences the magnitude of neuroinflammation and recovery trajectory, with APOE ε2 associated with reduced neutrophilic inflammation and shorter hospitalisation. Together, these findings identify ApoE-dependent and isoform-specific pathways as key contributors to inflammation resolution following encephalitis, warranting further mechanistic and translational investigation.

## Limitations of the study

Because recovery can only be identified retrospectively following reversal of weight loss, the earliest molecular events that commit animals to lethal versus non-lethal trajectories may precede the first timepoint analysed. The factors underlying divergent disease outcomes among genetically inbred female mice with comparable viremias remain unknown, but may include factors such as variation in microbiome composition^120^, social hierarchy and associated immunological effects^121,122^, or timing and magnitude of viral replication and inflammatory responses within circadian rhythms^123^. Rather than attempting to eliminate this biological variability, we exploited it to compare animals with divergent outcomes and thereby identify pathways specifically associated with neuroinflammation resolution. In addition, although the single-cell spatial transcriptomic dataset included a limited number of symptomatic recovered brains due to limited capacity on CosMx slides, the principal findings were independently supported by bulk RNA-seq, histopathology, genetic loss-of-function studies, and human clinical data. Finally, although our findings in acute encephalitis syndrome patients suggest that ApoE-dependent encephalitis resolution mechanisms may extend beyond Japanese encephalitis, whether these pathways are broadly conserved across neuroinvasive viral infections remains to be determined.

## Supporting information

Supplementary Table 12 - gsea_unique in recovered

Supplementary Table 13 - ipa_all

Supplementary Table 14 - ApoeKO and C57 bulk RNAseq raw differential expression data

Supplementary Table 15 - ApoeKO lethal vs C57 lethal pathway analyses IPA GO GSEA

Supplementary Table 1 - CosMx-Mouse-Neuroscience-Panel-Gene-Target-List

Supplementary Table 2 - CosMx JEV vs Mock pseudobulk

Supplementary Table 3 - CosMx recovered vs mock pseudobulk

Supplementary Table 4 - recovered vs JEV psueodbulk

Supplementary Table 5 - spatial psudobulk IPA

Supplementary Table 6 - Ligand-receptor analysis

Supplementary Table 7 - Apoe-Trem2-Tyrobp hotspot cells DE

Supplementary Table 8 - JEV virus read sequence changes

Supplementary Table 9 - Bulk RNAseq raw differential expression data

Supplementary Table 10 - Bulk RNAseq symptomatic recovered unique DEGs

Supplementary Table 11 - GSEA all raw data

Cox et al Supplementary Figures

## ACKNOWLEDGEMENTS

D.J.R. is an Emerging Leadership Fellow funded by the National Health and Medical Research Council (NHMRC) of Australia (Investigator Grant APP2041411). QIMR Berghofer received a generous philanthropic donation from the Brazil Family Foundation awarded to D.J.R. to support Japanese encephalitis virus research at QIMR Berghofer. Abigail Cox was a recipient of a PhD scholarship & fee waiver from the Faculty of Health, Medicine and Behavioural Sciences, University of Queensland, Brisbane, Australia.

From QIMR Berghofer we thank Dr I Anraku for managing the PC3 (BSL3) facility, animal house staff for mouse breeding and agistment, Crystal Chang and Sang-Hee Park for the histology and immunohistochemistry, and Tu Parsons and Paul Collins for the bulk RNA-seq.

## AUTHOR CONTRIBUTIONS

Conceptualisation, D.J.R. Methodology, D.J.R., C.R.B., A.L.C., and Q.H.N. Formal analysis, C.R.B., D.J.R., A.L.C., R.Z., G.H., M.P.H., Investigation, A.L.C., C.R.B., W.N., A.R.P., B.T., K.Y, and D.J.R. Resources, D.J.R., and A.S. Data curation, D.J.R., C.R.B., and A.L.C. Writing, original draft, D.J.R. and A.L.C. Writing, review and editing, C.R.B., A.L.C, and D.J.R. Visualisation, D.J.R., C.R.B., A.L.C., and G.H. Supervision, D.J.R., C.R.B., and W.N. Software, Q.H.N., R.Z., M.P.H., and C.R.B. Validation, G.H., C.R.B., and D.J.R. Project administration, D.J.R., W.N., and A.S. Funding acquisition, D.J.R, and A.S.

## DATA AVAILABILITY

All data is provided in the manuscript and accompanying supplementary files. Raw bulk RNA-seq data is available on the NCBI Sequence Read Archive with the bioproject accession PRJNA1481390. Raw single-cell spatial transcriptomics data are available through the Gene Expression Omnibus (GEO) under accession number GSE337780.

## Declaration of generative AI and AI-assisted technologies in the manuscript preparation process

During the preparation of this work the author(s) used M365 Copilot to enhance the clarity, language, and grammatical accuracy of the original text. After using this tool/service, the author(s) reviewed and edited the content as needed and take(s) full responsibility for the content of the published article.

