## Supplementary material for "Apolipoprotein E governs neuroinflammation resolution and recovery from viral encephalitis": Cox et al Supplementary Figures

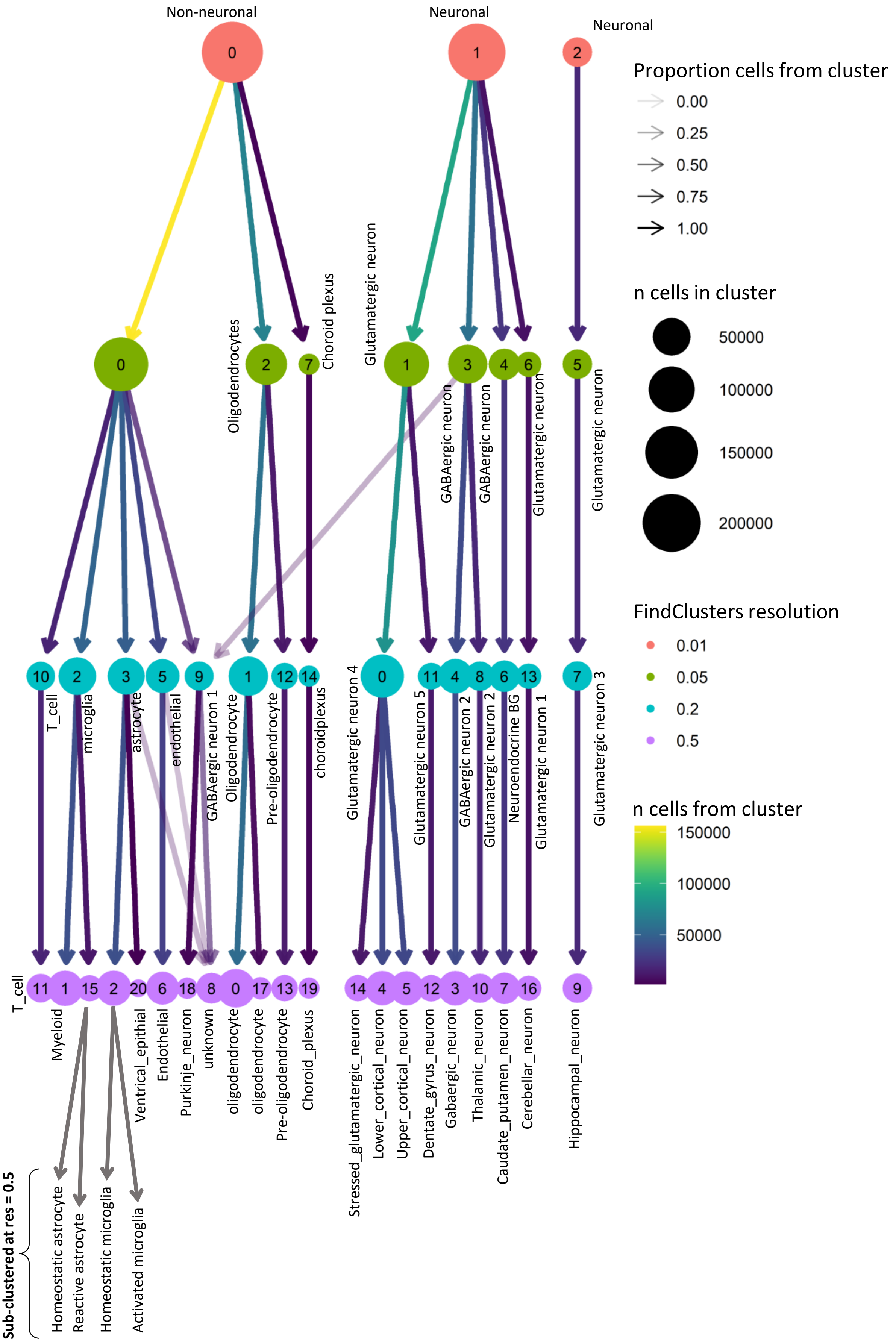


**Supplementary Figure 1.** Output from Clustree showing 4 levels of clustering resolution obtained using FindClusters. Clusters 2 and 15 were subjected to an additional round of clustering at resolution 0.5, yielding sub-clusters related to activation state, and resulting in a total of 22 annotated clusters.

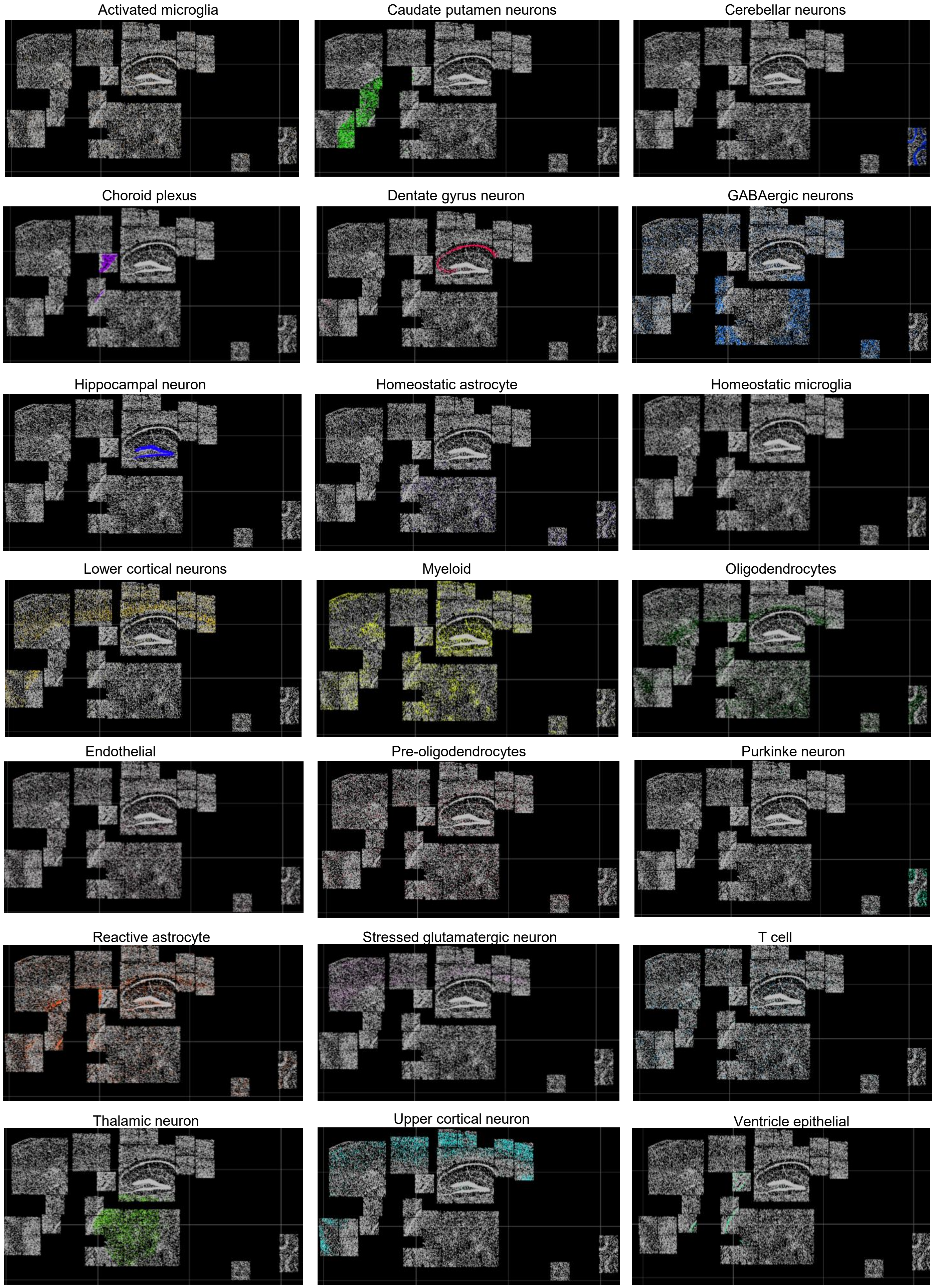


**Supplementary Figure 2**. A representative section showing cells coloured by annotated cluster.

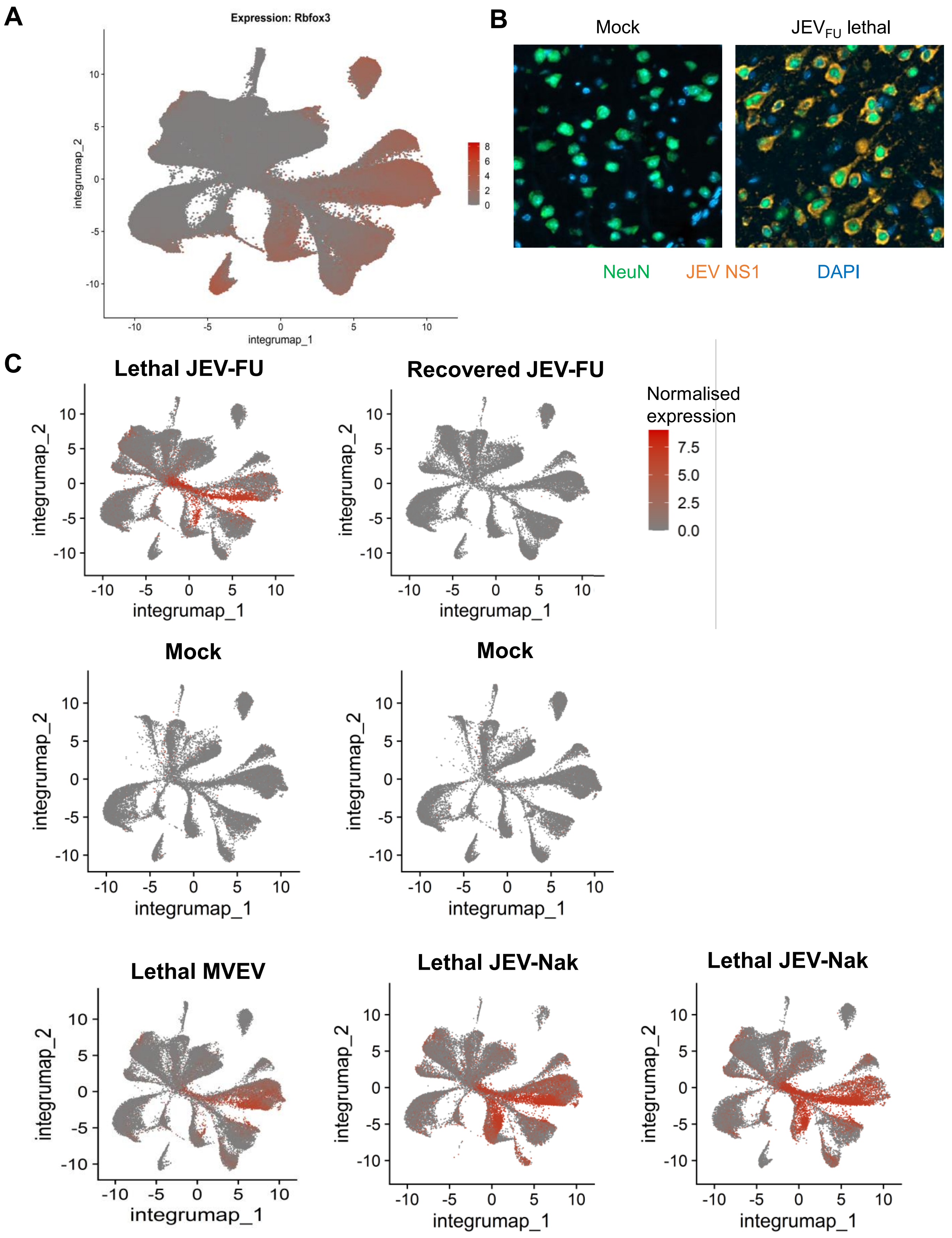


**Supplementary Figure 3.** A) Uniform Manifold Approximation and Projection (UMAP) of all cells in all experimental groups coloured according to normalised expression of Rbfox3. B) Sections from formalin fixed and paraffin embedded brains from mock or $JEV_{FU}$ infected (lethal outcome) brains from C57BL/6J mice were analysed by immunofluorescence. Sections were stained by indirect immunofluorescence with the anti-NS1 monoclonal antibody (4G4) (orange), and antibodies specific for a neuronal nuclear marker NeuN (green), with DAPI stain shown in blue. C) UMAP of each section coloured by viral probe expression.

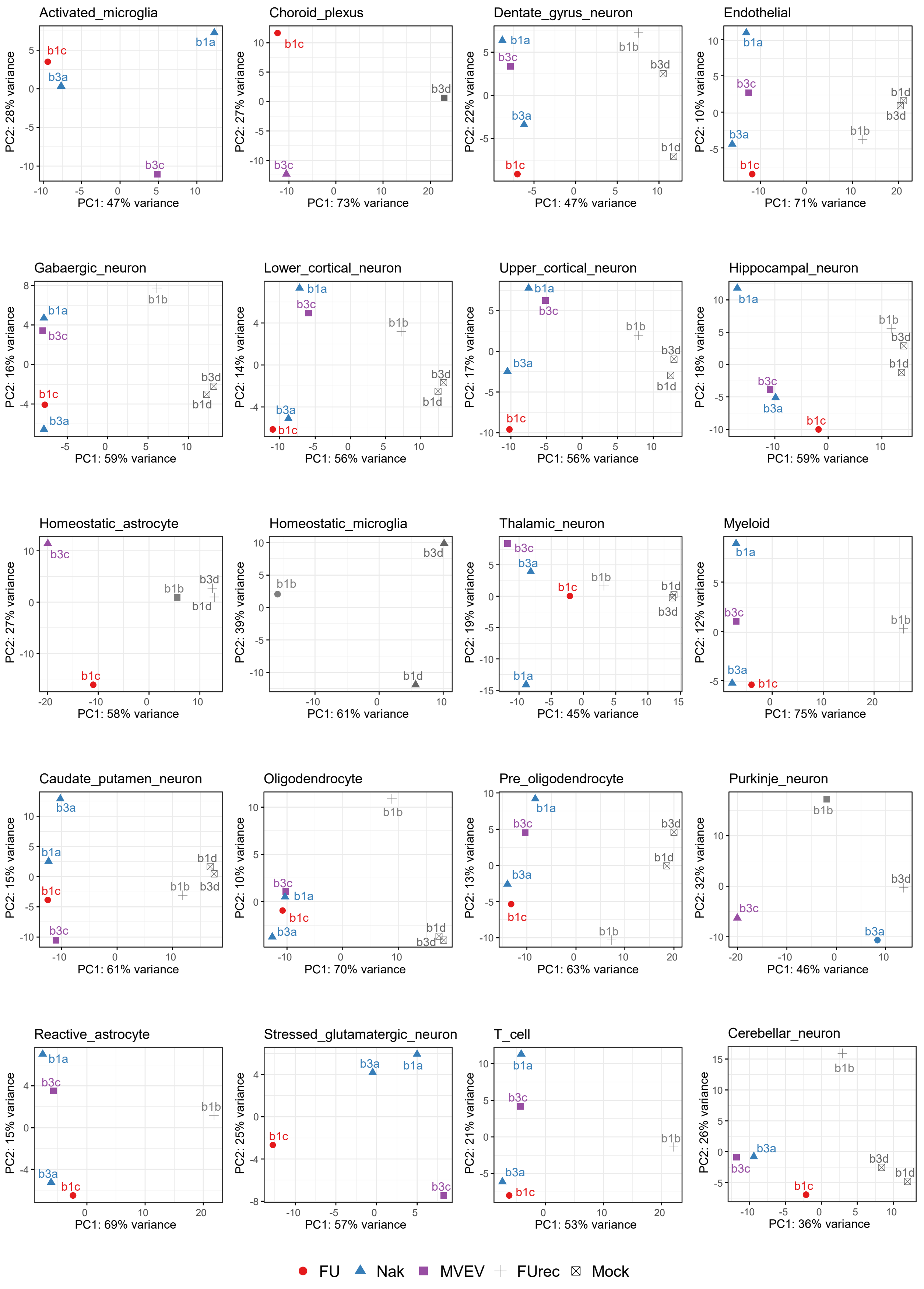


**Supplementary Figure 4.** Principle Components Analysis based on pseudo-bulk normalised gene expression profile per cell type. Each data point relates to a section, plotted by PC1 and PC2 values, and coloured by experimental group. FU = lethal JEV-FU; Nak = lethal JEV-Nak; MVEV = lethal MVEV; FUrec = recovered JEV-FU. Note that some cell types were absent in some sections or had counts below the pseudo-bulk QC threshold of ≥ 200. The Ventrical epithelial cell type was not included due to it only passing pseudo-bulk QC in 1 sample.

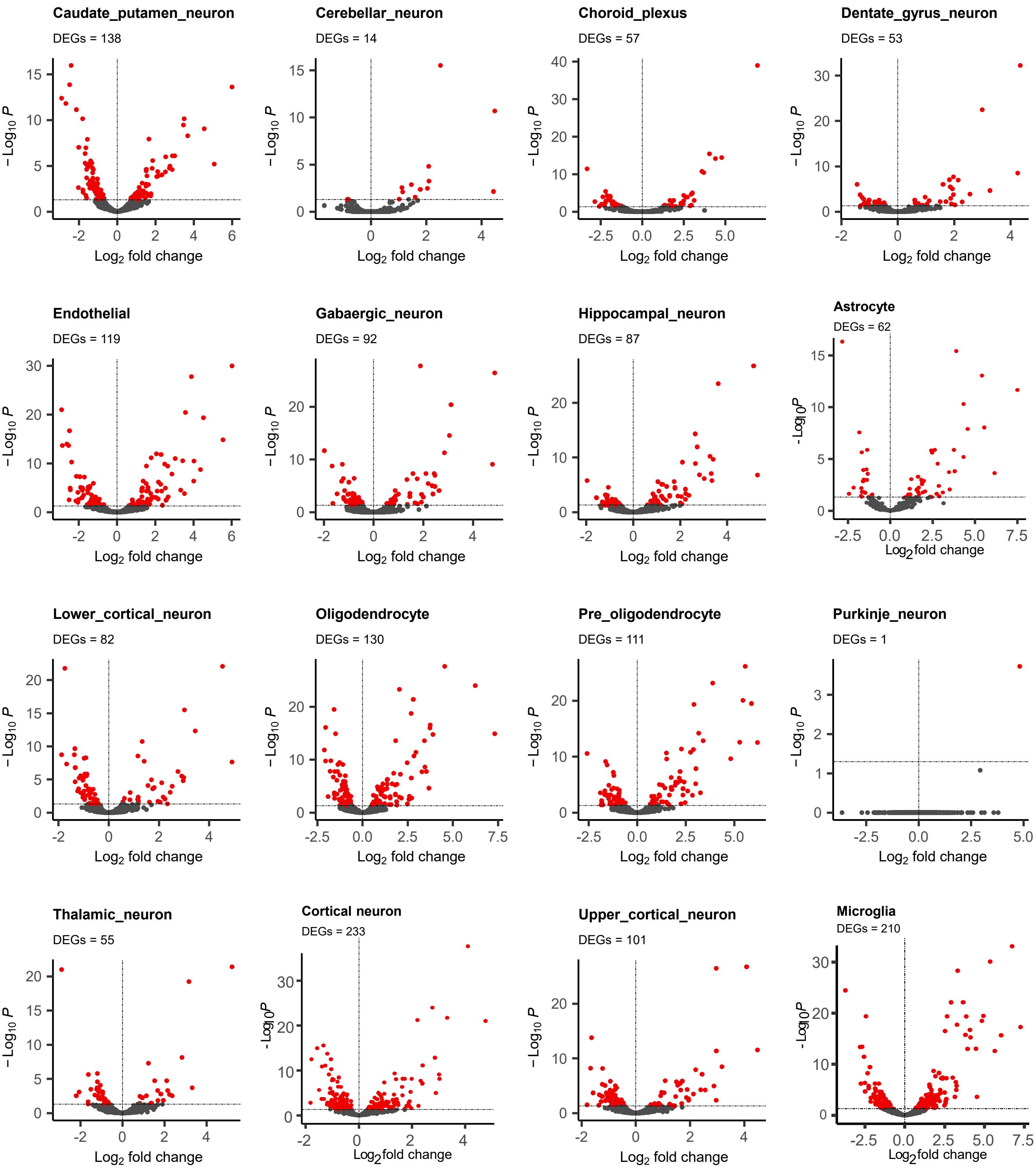


**Supplementary Figure 5.** Volcano plots show pseudo-bulk differential expression per cell type for lethal JEV versus mock groups. Y-axis is the negative $\log_{10}$ of the adjusted p-value according to DESeq2. Astrocytes represent the sum of reactive and homeostatic astrocytes. Microglia represent the sum of activated and homeostatic microglia. Cortical neuron represent the sum of upper- lower- and stressed-cortical neurons. Each point is one gene coloured according to significance: red = adjusted p-value < 0.05; black = not significant. Only those cell types that passed pseudo-bulk  QC and are present in each both conditions are shown.

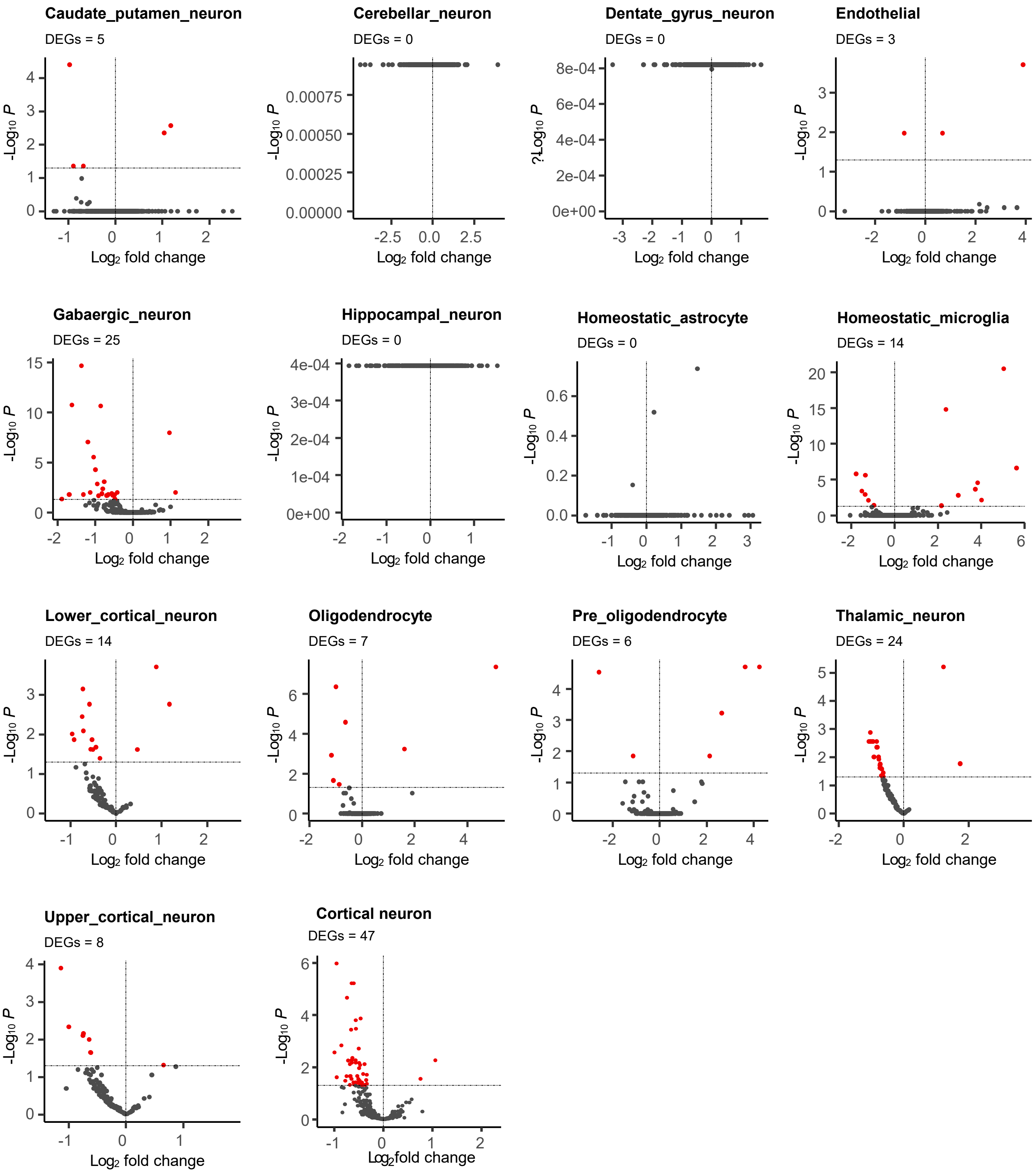


**Supplementary Figure 6**. Volcano plots show pseudo-bulk differential expression per cell type for recovered JEV versus mock groups. Y-axis is the negative $\log_{10}$ of the adjusted p-value according to DESeq2. Cortical neuron represents the sum of upper- lower- and stressed-cortical neurons. Each point is one gene coloured according to significance: red = adjusted p-value < 0.05; black = not significant. Only those cell types that passed pseudo-bulk QC and are present in each both conditions are shown.

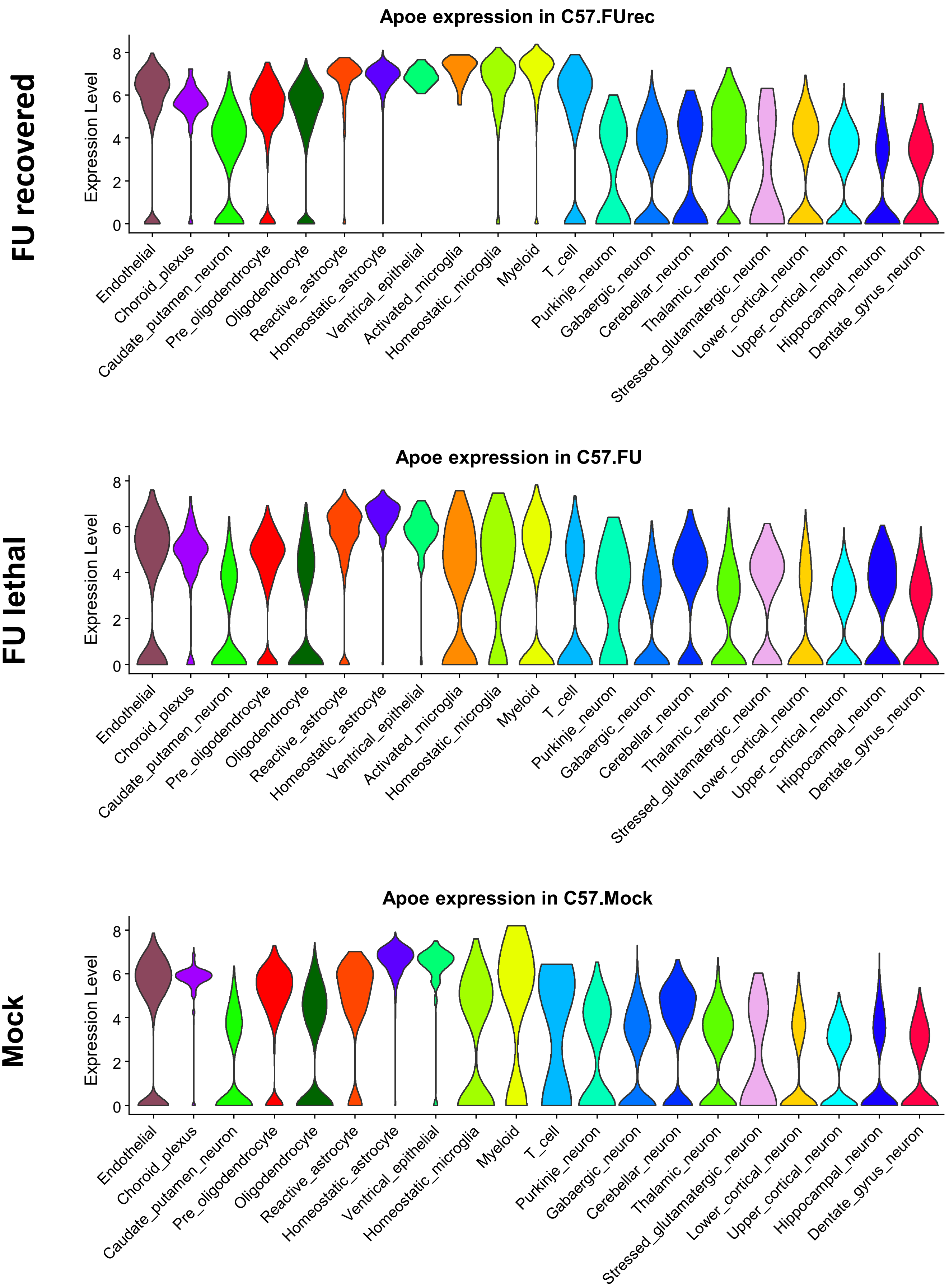


**Supplementary Figure 7.** Violins plots show distributions of per-cell Apoe normalised expression for all cell-types in each experimental group according to Seurat.

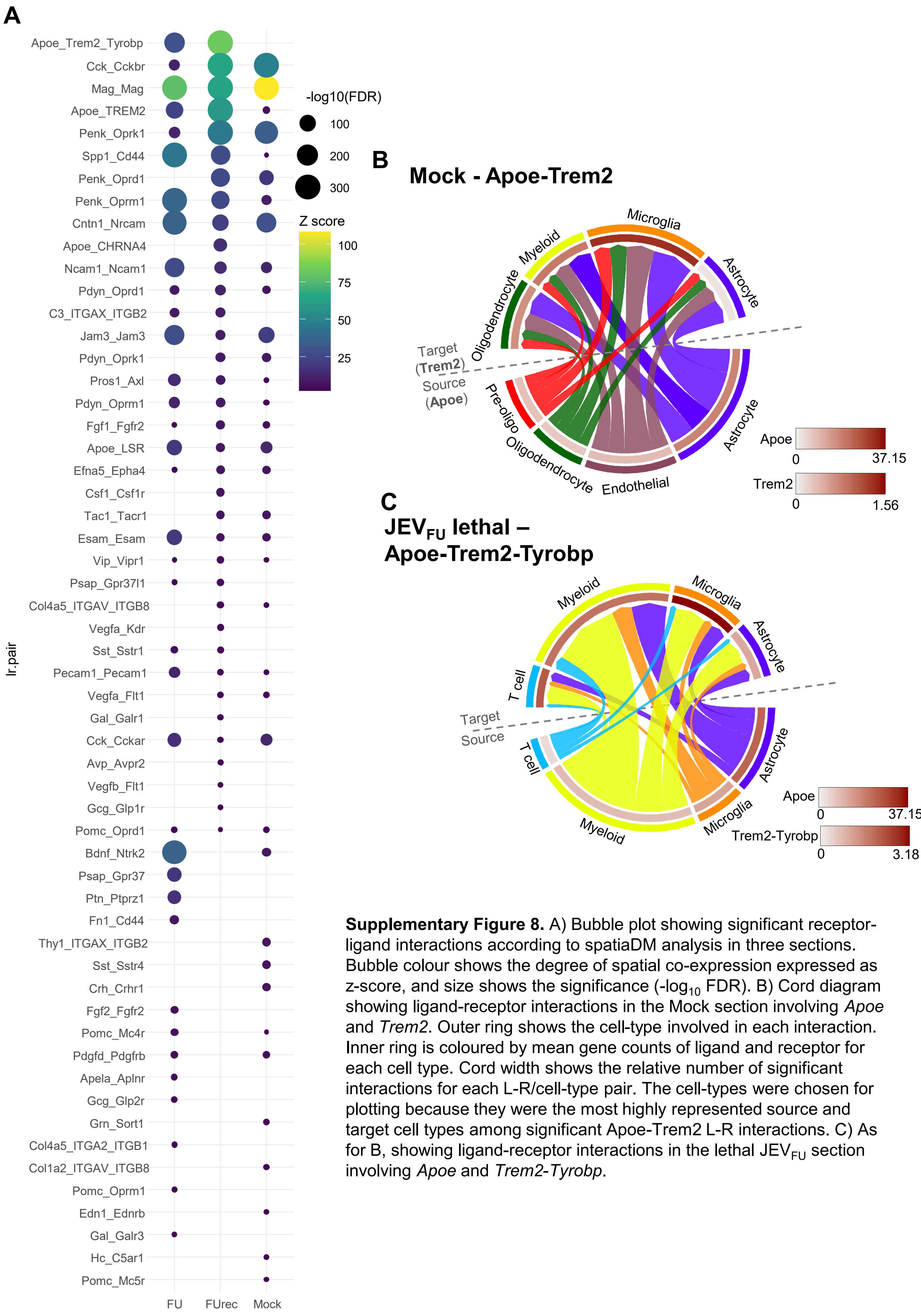


**Supplementary Figure 8.** A) Bubble plot showing significant receptor-ligand interactions according to spatiaDM analysis in three sections. Bubble colour shows the degree of spatial co-expression expressed as z-score, and size shows the significance (-log$_{10}$ FDR). B) Cord diagram showing ligand-receptor interactions in the Mock section involving *Apoe* and *Trem2*. Outer ring shows the cell-type involved in each interaction. Inner ring is coloured by mean gene counts of ligand and receptor for each cell type. Cord width shows the relative number of significant interactions for each L-R/cell-type pair. The cell-types were chosen for plotting because they were the most highly represented source and target cell types among significant Apoe-Trem2 L-R interactions. C) As for B, showing ligand-receptor interactions in the lethal JEV$_{FU}$ section involving *Apoe* and *Trem2-Tyrobp*.

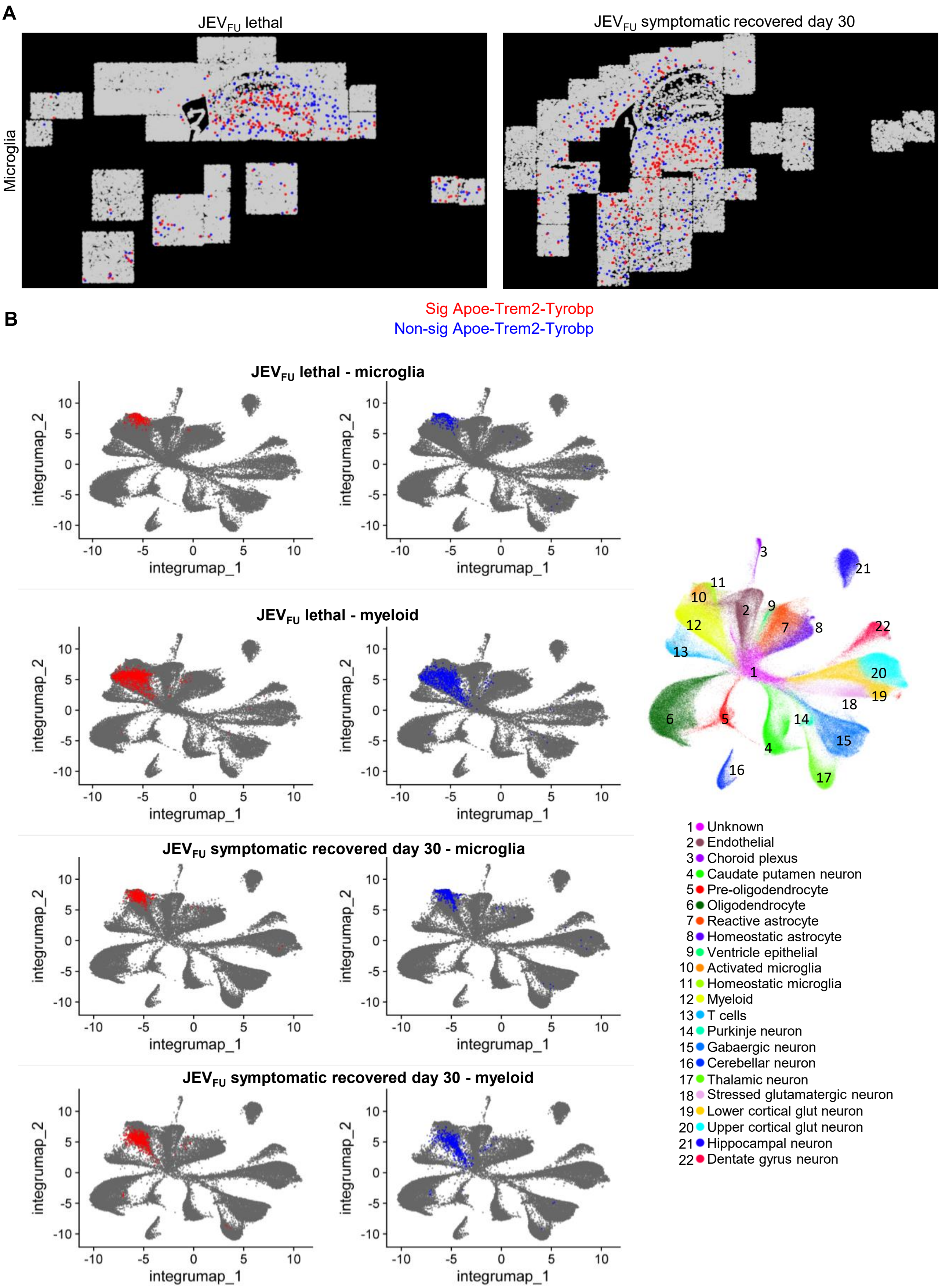


**Supplementary Figure 9. Microglia and myeloid cells that participate or not participate in *Apoe-Trem2-Tyrobp* ligand-receptor interactions.** A) Spatial distribution of 'Apoe-receiver' microglia with significant ($p<0.05$) SpatialDM and receptor Moran's I > 1 (significant, red), and 'Apoe-non-receiver' cells (non-significant, blue), for a JEV symptomatic recovered (right) and lethal (left) section. B) The significant (red) and non-significant (blue) cells from 'A' and Figure 3G shown on UMAP plots. Cell clustering from Figure 1A is shown for reference.

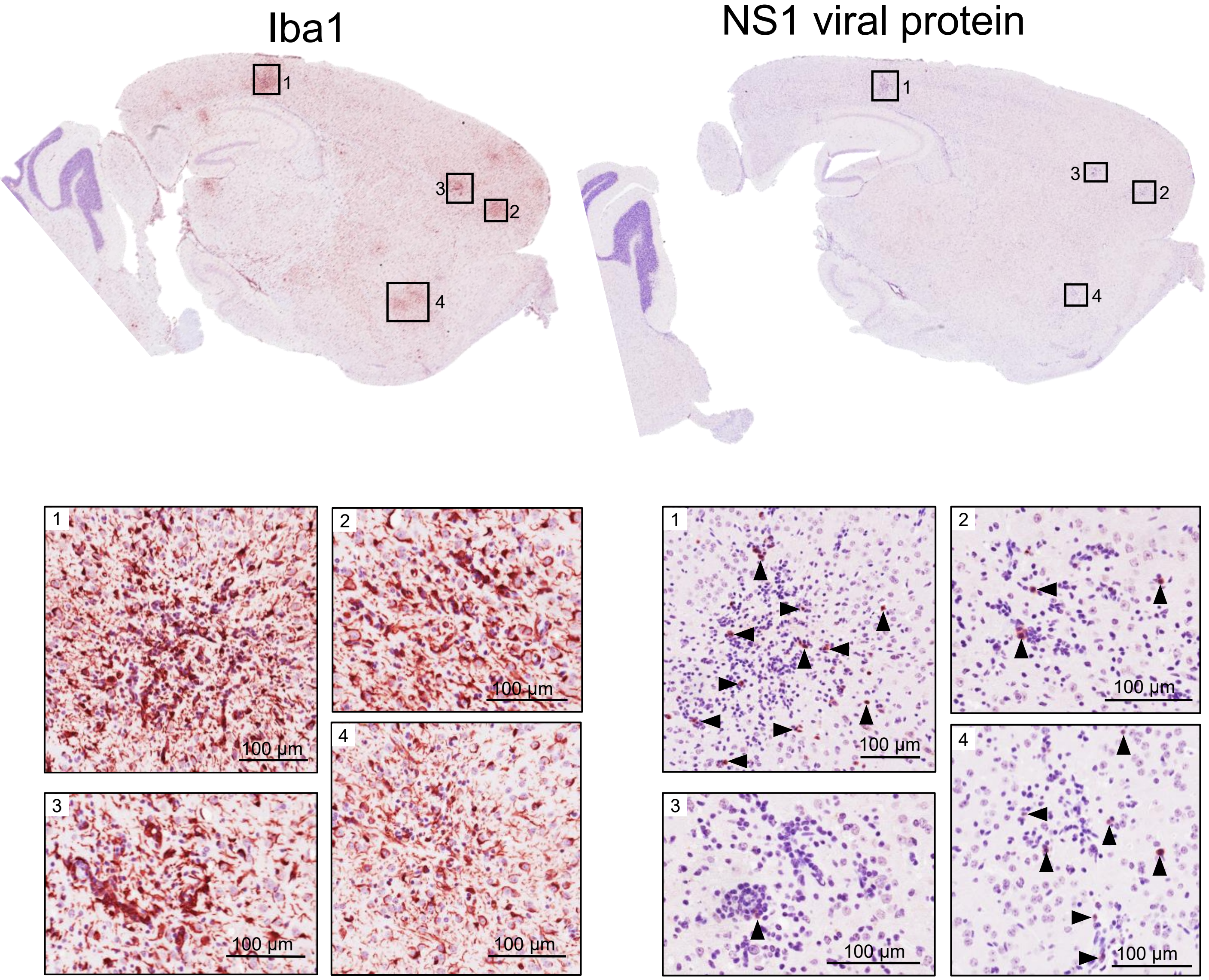


**Supplementary Figure 10.** In brains from symptomatic recovered mice, focal regions of pronounced microglial and myeloid activation, co-localised with cells expressing JEV NS1 protein. Immunohistochemistry performed with an Iba1 antibody (left) for myeloid and microglial cells, and an NS1 antibody (4G4; right) for viral infected cells is shown for the same brain section. Insets labelled 1-4 correspond to the indicated regions and highlight identical areas across the two stains.

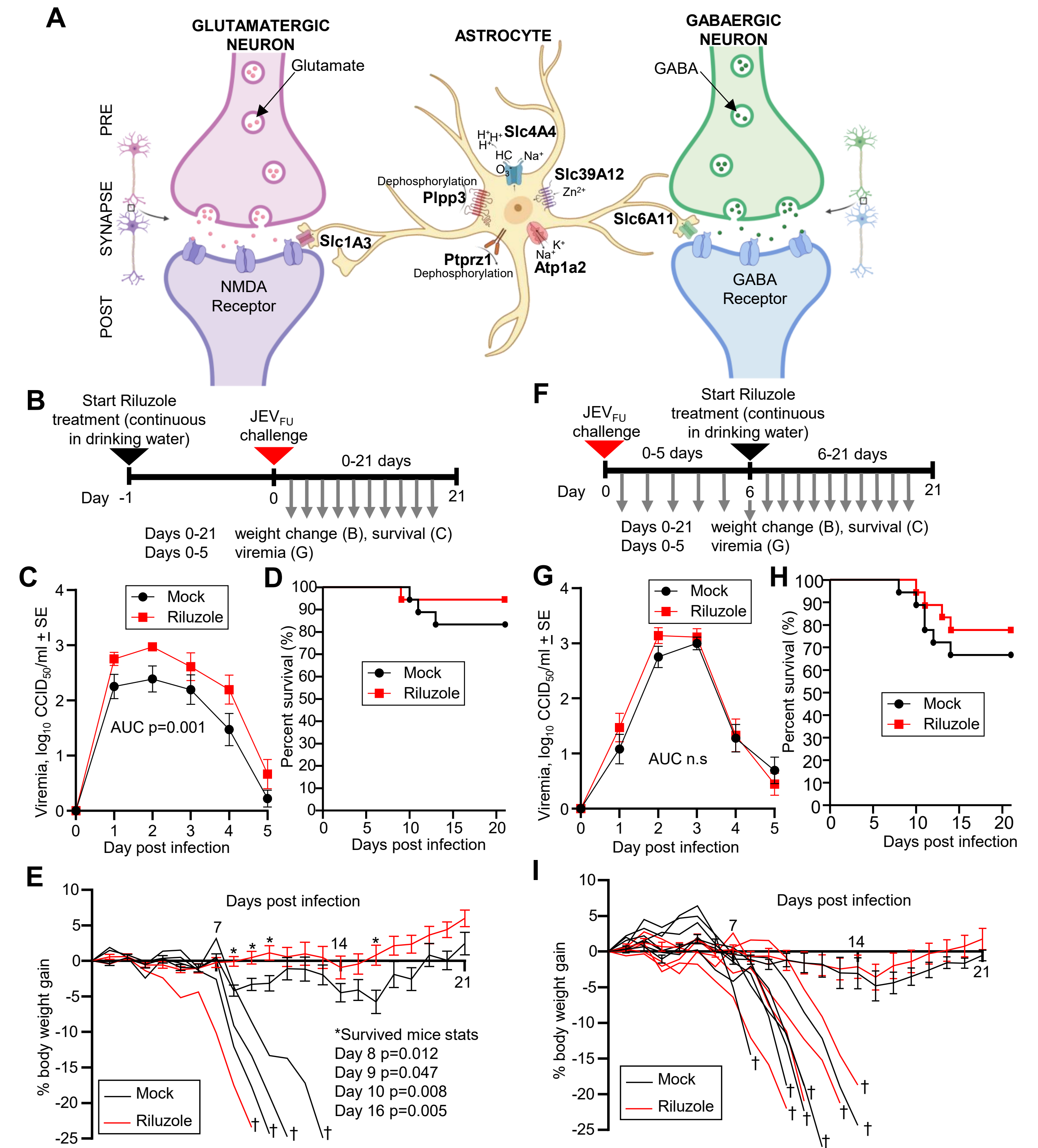


**Supplementary Figure 11. A)** Schematic overview of astrocyte- and glia-associated genes involved in neurotransmitter clearance, ion and pH homeostasis, lipid signalling, and myelin regulation, including Slc1a3 (EAAT1-mediated glutamate uptake), Slc6a11 (GABA uptake), Atp1a2 (Na+/K+ homeostasis), Slc4a4 (bicarbonate-dependent pH regulation), Slc39a12 (zinc import), Plpp3 (inactivation of pro-inflammatory lipids), and Ptprz1 (oligodendrocyte-associated myelination). **B)** Female ≈6-week-old C57BL/6J mice were infected s.c. with $5 \times 10^3$ $CCID_{50}$ of $JEV_{FU}$. **B) and F)** Timeline of experiments. Silver arrows indicate sample/data collection timepoints. **C), G)** Viremia comparing Riluzole-treated mice (red, n=18) with mock-treated mice (black, n=) over 5 days. Limit of detection is 2 $log_{10}CCID_{50}$/ml. Statistics represent t-test or Kolmogorov-Smirnov exact test for area under the curve (AUC) values. **D), H)** Kaplan-Meier plot showing percent survival for Riluzole-treated mice (red, n = 18) and mock-treated mice (black, n = 18). **E)** Percent body weight change of individual mice with lethal (†) outcomes after JEV infection for Riluzole-treated mice (red, n = 1) and mock-treated mice (black, n = 3). For the surviving mice from each group, their body weights were averaged for each day. (Riluzole-treated mice (red, n = 17), mock-treated mice (black, n = 15)). Statistics represent t-test or Kolmogorov–Smirnov exact test at the indicated timepoints. **I)** Percent body weight change of individual mice with lethal (†) outcomes after JEV infection for Riluzole-treated mice (red, n = 4) and mock-treated mice (black, n = 6). For the surviving mice from each group, their body weights were averaged for each day. (Riluzole-treated mice (red, n = 14), mock-treated mice (black, n = 12)).

| | One day post peak weight loss | Symptomatic day 21 | Symptomatic day 30 |
|---|---|---|---|
| Phagocytosis and cross presentation | Xcr1 lfc 4.41, fdr 0.002<br>MM_GOBP_IMMUNE_RESPONSE_REGULATING_CELL_SURFACE_RECEPTOR_SIGNALING_PATHWAY_INVOLVED_IN_PHAGOCYTOSIS, NES 1.63, fdr 0.016<br>Phagosome formation, z-score 0.82, -log(p-value) 0.95 | Xcr1 lfc 4.57, fdr 0.003<br>Communication between Innate and Adaptive Immune Cells, z-score 2.2, -log(p-value) 3.46<br>Phagosome formation, z-score 2, log(p-value) 3.14 | |
| M2 macrophage | Rgs10 lfc 0.65, fdr 0.004<br>Arhgap15 lfc 0.87, fdr 0.047 | Rgs10 lfc 0.79, fdr 0.0005<br>**Apoe lfc 0.89 fdr 0.005**<br>IL-12 Signaling and Production in Macrophages, z-score -2, -log(p-value) 3.95<br>TGFB, z-score 0.7, p-value 0.0002 | ***Apoe lfc 1.06 fdr 0.006 vs lethal*** |
| Treg and Th2 | Itgae lfc 2.00, fdr 0.001<br>Ccl22 lfc 2.23, fdr 0.01<br>MM_ZHENG_FOXP3_TARGETS_IN_THYMUS_UP, NES 1.63, fdr 0.15<br>TCR (complex), z-score 2.6, p-value 0.002<br>IL5, z-score 1.96, p-value 0.046 | Zfp683 lfc 5.59, fdr 0.005<br>Itgae lfc 3.88, fdr 2E-12<br>Cd4 lfc 0.7, fdr 0.03<br>MM_ZHENG_FOXP3_TARGETS_IN_THYMUS_UP, NES 1.63, fdr 0.15<br>TCR (complex), z-score 2.6, p-value 0.002<br>IL5, z-score 1.94, p-value 0.0002 | |
| Plasma cells and antibody responses | Ighm lfc 1.4, fdr 0.002 | Ighm lfc 3.25, fdr 2.04E-15<br>Jchain lfc 6.33, fdr 2.00E-12<br>MM_MORI_PLASMA_CELL_UP, NES 1.6, fdr 0.009<br>Immunoglobulin (complex), z-score 1.25, p-value 0.0000012 | Ighm lfc 2.75, fdr 0.038<br>Jchain lfc 6.42, fdr 0.006 |
| Lipid metabolism | Fabp7 lfc 1.11, fdr 0.044<br>Elovl5 lfc 0.33, fdr 0.003<br>MM_GOBP_CHOLESTEROL_EFFLUX, NES 1.57, fdr 0.036<br>Eicosanoid signalling, z-score -1.34, -log(p-value) 2<br>MLXIPL, z-score 3, p-value 4.64E-08 | **Apoe lfc 0.89 fdr 0.005**<br>Abca9 lfc 0.81, fdr 3.3E-06<br>**MM_GOMF_APOLIPOPROTEIN_BINDING, NES 1.58, fdr 0.03**<br>MM_REACTOME_CHOLESTEROL_BIOSYNTHESIS, NES -2.05, fdr 0.004<br>Fatty acid metabolism, z-score 1, p-value 0.0005 | ***Apoe lfc 1.06 fdr 0.006 vs lethal***<br>**MM_GOMF_APOLIPOPROTEIN_BINDING, NES 1.73, fdr 0.009**<br>MM_REACTOME_CHOLESTEROL_BIOSYNTHESIS, NES -1.85, fdr 0.0495 |
| Tissue repair | MYC, z-score 4.04, p-value 0.0000003<br>Cell cycle progression, z-score 1.18, fdr 0.005<br>Necrosis, z-score -2.45, p-value 0.0005 | Thbs4 lfc 0.91, fdr 0.002<br>Organismal death, z-score -1.43, p-value 0.0001 | Slc1a3, lfc 1.32, fdr 0.003<br>Ptprz1 lfc 1.12, fdr 0.01<br>Slc39a12 lfc 1.06, fdr 0.049<br>Slc4a4 lfc 1.04, fdr 5.4E-05<br>MM_LEIN_ASTROCYTE_MARKERS, NES 1.7, fdr 0.01<br>MM_GOMF_COLLAGEN_BINDING, NES 1.85, fdr 0.0008<br>Cell viability, z-score 2, p-value 0.02<br>Organismal death, z-score -1.91, p-value 0.008 |

Unique DEGs　　Unique GSEAs

IPA USRs from unique DEGs　　IPA Canonical pathways from unique DEGs　　IPA Diseases and functions from unique DEGs

**Supplementary Figure 12**. Summary of bulk RNA-seq analyses highlighting differentially expressed genes (DEGs) and enriched pathways uniquely associated with symptomatic recovery and absent in lethal disease. Data are shown for three timepoints: one day post-peak weight loss (left), symptomatic day 21 (middle), and symptomatic day 30 (right). Shown are common biological processes identified across these timepoints, including phagocytosis and antigen cross-presentation, M2 macrophage polarisation, T regulatory (Treg) and Th2 responses, plasma cell and antibody responses, lipid metabolism, and tissue repair. Different data and analysis types are indicated by colour: unique DEGs (black), unique GSEA gene sets (red), IPA upstream regulators derived from unique DEGs (green), IPA canonical pathways derived from unique DEGs (purple), and IPA diseases and functions derived from unique DEGs (blue). Data were selected from raw analysis outputs provided in Supplementary Tables 10, 12, and 13.

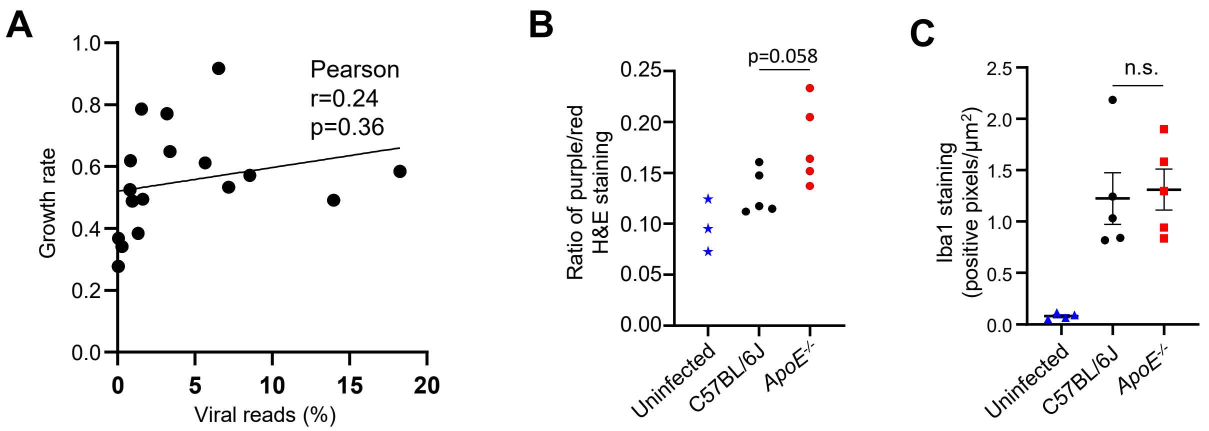


**Supplementary Figure 13.** A) Pearson correlation for viral RNA reads as a percentage of total reads (from Figure 5A) and growth rate (calculated from Figure 6C). B) Ratio of nuclear (dark purple) to non-nuclear (red) staining of H&E stained brain sections (a measure of leukocyte infiltration). Individual mice brains are shown for uninfected (blue stars, n=3), C57BL/6J JEV infected with lethal outcomes (black circles n=5), and *Apoe$^{-/-}$* JEV infected with lethal outcomes (red circles, n=5). Data are the same lethal mice as shown in Figure 6D-L. Statistics by t-test. C) Quantification of Iba1 staining (positive pixels/µm$^2$) in the same brains as for ‘B’. Statistics by t-test.

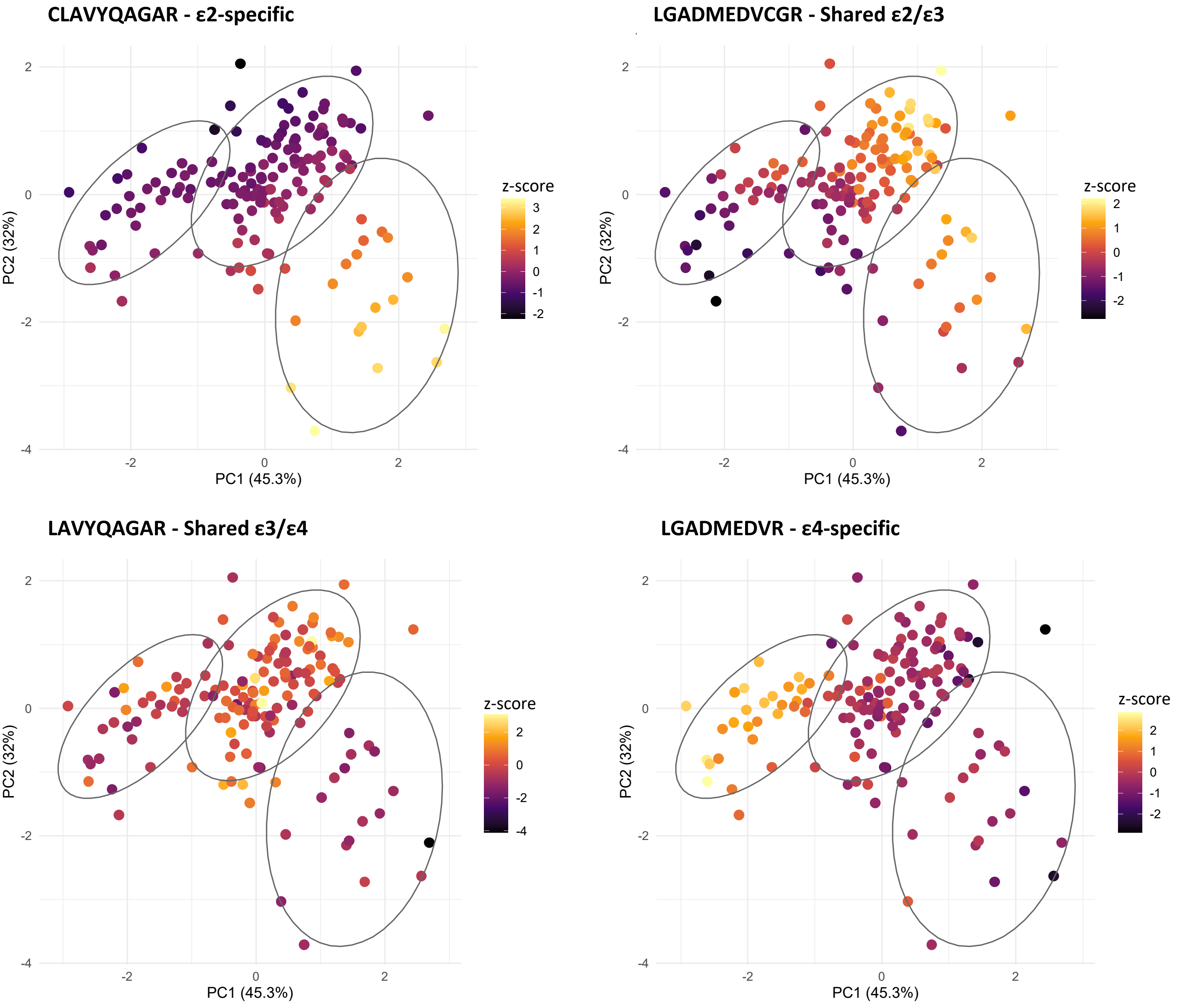


**Supplementary Figure 14.** Principle Components Analysis of 148 acute encephalitis samples using normalised abundances of three APOE isoform-associated peptides: CLAVYQAGAR (ε2-specific), LGADMEDVCGR (Shared ε2/ε3), and LGADMEDVR (ε4-specific). This combination of peptides was chosen for 3D-GMM clustering of samples for genotyping. Points are coloured by expression of each of the four APOE isoform-associated peptides (z-score).
